# Flippable Siamese Differential Neural Network for Differential Graph Inference

**DOI:** 10.1101/2025.08.31.673390

**Authors:** Jiacheng Leng, Jiating Yu, Ling-Yun Wu

## Abstract

Differential graph inference is a critical analytical technique that enables researchers to accurately identify the variables and their interactions that change under different conditions. By comparing two conditions, researchers can gain a deeper understanding of the differences between them. Currently, the mainstream methods in differential graph inference are mathematical optimization algorithms, including sparse optimization based on Gaussian graphical models or sparse Bayesian regression. These methods can eliminate many false positives in graphs, but at the cost of heavy reliance on the prior distributions of data or parameters, and they suffer from the curse of dimensionality. To address these challenges, we introduce a new architecture called the Flippable Siamese Differential Neural Network (FSDiffNet). We originally established the concept of flippability and the theoretical foundation of flippable neural networks, laying the groundwork for building a flippable neural network. This theoretical framework guided the design of architecture and components, including the SoftSparse activation function and high-dilation circular padding diagonal convolution. FSDiffNet uses large-scale pre-training techniques to acquire differential features and perform differential graph inference. Through experiments with simulated and real datasets, FSDiffNet outperforms existing state-of-the-art methods on multiple metrics, effectively inferring key differential factors related to conditions such as autism and breast cancer. This proves the effectiveness of FSDiffNet as a solution for differential graph inference challenges.

## Introduction

Graph inference is a key area in graph theory and machine learning, aiding in the analysis of interactions between variables from observed data. In this paper, we focus on Markov Random Fields (MRF), which precisely describe complex variable interactions through a crucial characteristic: the weight *W*_*ij*_ is zero only if variables *i* and *j* are conditionally independent given the rest of the variable set *V* \ {*i,j*}.

However, inferring graphs under a single condition often fails to reveal significant variations in the relationships between variables under different conditions. For example, in genomics, differential graph inference identifies driver genes by inferring gene regulatory networks from normal and cancerous samples, thereby enhancing our understanding of the factors driving diseases^1–3^.

In recent years, there has been a growing interest within the scientific community in the study of differential graph inference. This area of research can be primarily divided into two main approaches. The first leverages Maximum Likelihood Estimation (MLE) within Gaussian Graphical Models (GGM). ^4–10^. This technique involves tackling optimization problems based on the assumption of Gaussian distributions, an assumption that often does not align with the complexities of real-world scenarios. Alternatively, another strand of research focuses on Bayesian graph learning through the use of Markov Chain Monte Carlo (MCMC) sampling methods ^11–14^. While this approach provides a comprehensive framework, it is also characterized by high computational demands due to the extensive nature of the sampling processes involved. Parallel to these developments, there has been significant progress in applying deep learning techniques to graph inference ^15–22^. However, the creation of architectures specifically designed for differential graph inference remains relatively rare. One of the major challenges in this area is ensuring input flippability within the models. This means that the output of the model should change inversely when the order of inputs is reversed, a requirement that poses significant challenges, especially within Convolutional Neural Network (CNN) frameworks. Moreover, considering every possible combination of variables would result in overwhelming computational complexity. Therefore, most current models focus on predicting a single interaction between two variables at a time, treating the issue as a binary classification problem of determining whether an interaction exists. However, this method often neglects the impact of other variables on the variables of interest, potentially leading to biased results and the loss of vital information.

To tackle the challenges in differential graph inference, we introduce a novel architecture: the Flippable Siamese Differential Neural Network (FSDiffNet). This network innovatively incorporates a diagonal convolutional kernel (DiagConv) for comprehensive element coverage, essential for graph inference learning. Additionally, we develop a new SoftSparse activation function to effectively address the flippability issue, ensuring model outputs inversely change with the input order. Central to this work is the concept of flippable mapping, from which we derive the necessary conditions for designing neural networks suitable for differential graph inference. FSDiffNet utilizes siamese layers subtraction to facilitate differential graph inference, marking a significant advancement in this field.

Besides the special novel structure, we also demonstrate several features of FSDiffNet that facilitate flexible use, such as input flexibility, permutability of nodes, scalability of nodes numbers and sample numbers, and the non-specificity of the distributions. Finally, the experiments on both simulation datasets and real-world datasets suggest that FSDiffNet outperforms other state-of-the-art methods in several metrics, which validate that FSDiffNet is a useful architecture for differential graph inference problems.

Our work presents several key contributions in the field of neural network architecture design, particularly focusing on differential graph inference. We address the challenge of unbalanced receptive fields by employing a novel approach that combines circular padding with high dilation, ensuring a more balanced field of perception across the graph adjacency matrix. The cornerstone of our research is the establishment of a series of theories on flippable mapping, which aid in the design of neural network architectures for differential graph inference, namely FSDiffNet. To verify the effectiveness of FSDiffNet, we conducted extensive experiments on various datasets, including extreme challenge scenarios and real-world datasets such as autism and TCGA breast cancer datasets. These experiments consistently demonstrated the robust performance of FSDiffNet and its broad application potential in the field.

## Results

### Overview of FSDiffNet

The workflow of FSDiffNet is shown in Figure 1. FSDiffNet’s main task is to learn the inference of differential graphs. The data flow during the training process is illustrated in Fig. 1a. Initially, we generate a large number of graphs with random structures and establish differential patterns to derive graph pairs. Subsequently, utilizing Copula Gaussian graphical models, we simulate a vast array of observational data characterized by complex mixed distributions based on the precision matrices constructed from these graph pairs. Finally, we compute the correlation coefficient matrices from this observational data to serve as inputs for our model. We use the known graph structures as targets to train a differential graph inferencer, which can be used for efficiently and rapidly inferring differential graphs on real data, assisting researchers in conducting the studies based on differential interactions for the downstream tasks.

**Fig. 1:**
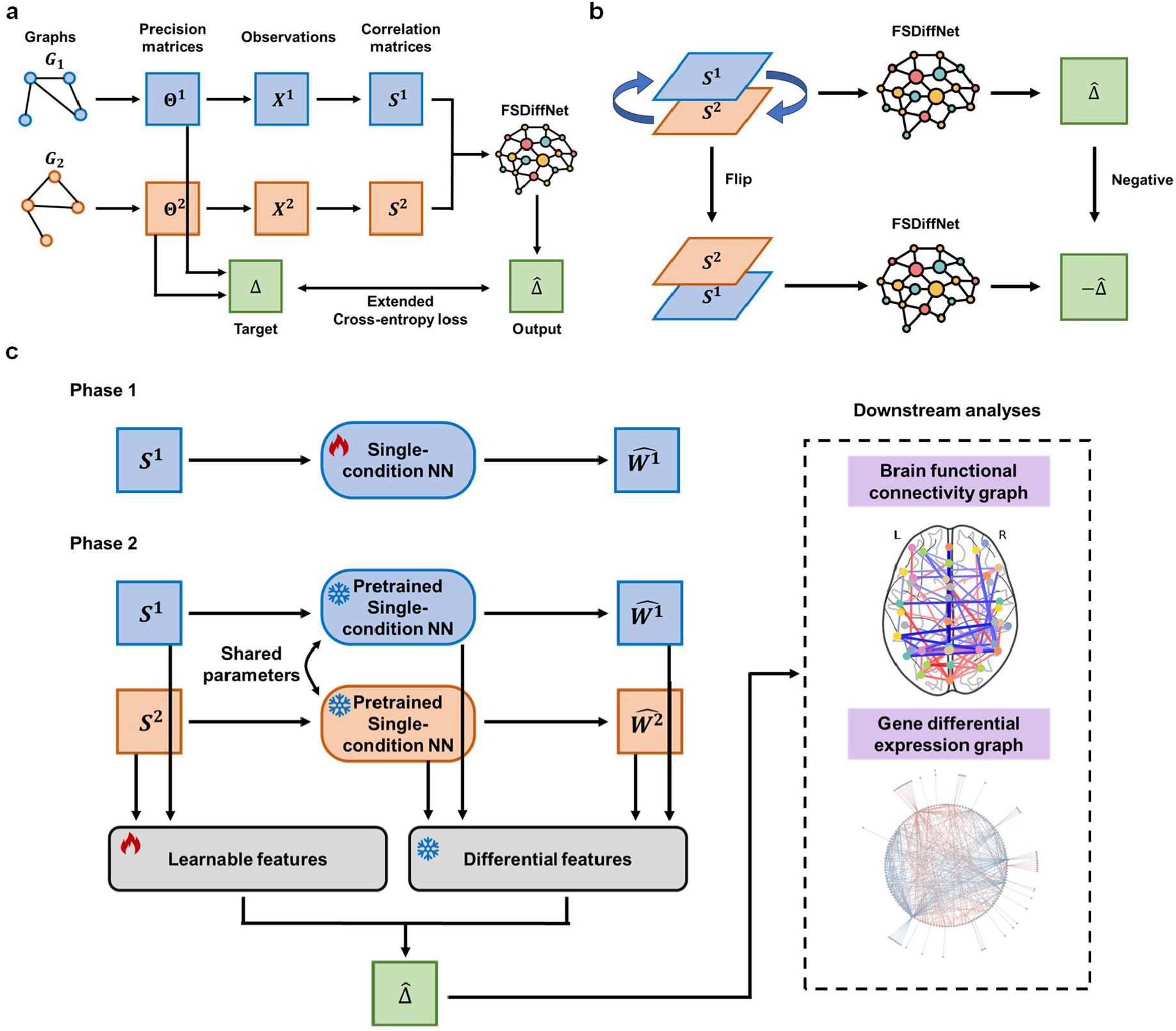
The workflow of FSDiffNet. **a** The data flow during the training process of FSDiffNet. **b** The flippability of FSDiffNet. When the input channels flip, the output strictly becomes opposite. **c** The overall training process of FSDiffNet. Phase 1 involves the pre-training of neural networks for single-condition graph inference. Phase 2 involves re-learning for differential graph inference tasks by constructing siamese networks with pretrained parameters in Phase 1. The flames represent the parameters and features that are learnable, whereas the snowflakes signify those are fixed.

The input of FSDiffNet consists of two correlation coefficient matrices with diagonals of 1 under two different conditions, and the output is an inferred undirected graph’s adjacency matrix. This matrix is weighted and signed, representing the differential graph, which includes potential key factors driving the differences. An important and novel feature is that FSDiffNet can obtain completely opposite results when the input channels flip, which is quite rare in neural network frameworks (Fig. 1b).

FSDiffNet’s training comprises two stages (Fig. 1c). The first phase is the pre-training phase of the single-condition baseline model. The input of the single-condition baseline model is a correlation coefficient matrix, and the output is an inferred undirected graph adjacency matrix, which is also weighted and signed, representing the single-condition graph structure. The second phase is the continual training of the two-condition flippable siamese neural network. The siamese neural network is a coupled architecture based on two artificial neural networks. It consists of two subnetworks with identical structures and shared weights. We input the correlation coefficient matrices from two different conditions into the respective sub-networks of the siamese neural network, which yields the outputs and features for each sub-network individually. By subtracting these features, we obtain differential features, which do not require additional training to generate. To capture more dynamic differential information, we input the subtracted input data into a learnable channel, from which we derive the learnable features. These features, combined with the frozen features, are then coupled through a 1×1 convolution layer to produce the final output.

Most importantly, we have established a theoretical framework for flippability (see Methods), which guides our architectural design of FSDiffNet. This theory ensures that the neural network maintains flippability in processing inputs, meaning that 𝒢 (*X*^1^,*X*^2^) = −𝒢(*X*^2^,*X*^1^) While implementing flippability is relatively straightforward in manually designed differential graph inferencers, it poses greater complexity in neural networks. Common approaches, such as subtracting results from a single inferencer, tend to lose crucial coupling information between conditions. To preserve flippability through neural layers and facilitate joint learning, our architecture incorporates specific design elements, such as odd convolutional layers and SoftSparse activation functions. These elements are grounded in our theoretical insights into the decomposability of flippable mappings and the conditions necessary for designing flippable neural networks, enabling FSDiffNet to effectively handle input order and coupled data (see Methods).

### Architecture and components of FSDiffNet guarantee correctness and performance of differential graph inferrence

The network architecture of FSDiffNet, as shown in Figure 2a, comprises three parallel channels. The first two channels are part of a siamese neural network sharing parameters, obtained through pretraining on single-condition graph inference tasks. The third channel integrates differential and learnable features, serving as the training channel for differential graph inference. We also designed network components such as diagonal convolution layers with high dilation rates and circular padding, SoftSparse activation function layers, and odd batch normalization layers.

**Fig. 2:**
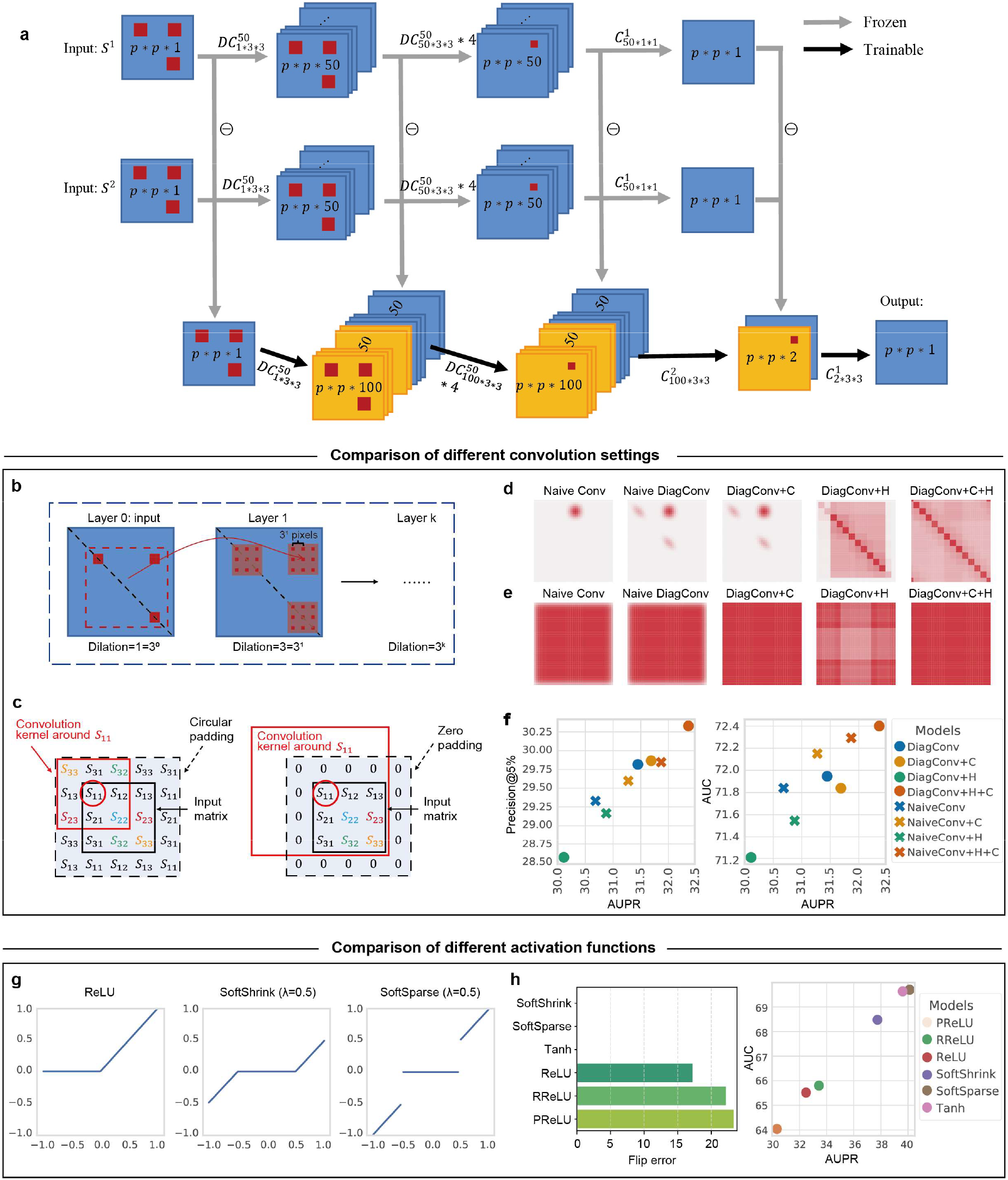
FSDiffNet and its components. **a**.The network architecture of FSDiffNet, where grey arrows indicate pre-trained parameters. Black arrows represent parameters that need learning. Blue channels are deterministic, while yellow channels are learnable. 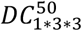 denotes a diagonal convolution layer with an input channel count of 1 and an output channel count of 50, using a 3×3 convolution kernel positioned at the top-right block. 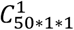 represents denotes a ordinary convolution layer with an input channel count of 1 and an output channel count of 50, using a 1×1 convolution kernel. ⊝ represents the subtraction. *p* is the number of variable. The receptive field of diagonal convolution at each layer. **c**. A comparison between circular padding and zero padding. **d**. The receptive field of element *Y*_5,20_ under different convolution settings. **e**. The distribution of receptive fields for all elements under different convolution settings. **f**. Performance comparison under different convolution settings. The horizontal axis represents the average AUPR, while the vertical axis shows average precision@5% and AUC. **g**. Comparison of different activation functions. **h**. Comparison of flip errors and performance for different activation functions.

Here, we introduce diagonal convolution (DiagConv), which establishes intrinsic connections between nodes. Unlike traditional convolution kernels that cover the area {*a,b,c*}×{*d,e,f*} (row×column), DiagConv also covers two diagonal sections {*a,b,c*}× {*a,b,c*} and {*d,e,f*}× {*d,e,f*} (where *a,b,c* or *d,e,f* need not be consecutive), as shown in the top right red block area of Figure 2b, layer one. This design allows the convolution kernel to encompass interactions within a set of points, such as {*a,b,c*}× {*a,b,c*}.and with increasing layers, this kernel can quickly cover all elements. Since graph inference requires all entries of the input matrices, we introduced a high dilation rate to expand the receptive field with as few parameters and layers as possible. To design a more efficient graph inferencer, we also combined circular padding. As our goal is to infer graph structures (a variant of inverse covariance matrices), it is necessary to utilize information from algebraic cofactors, and circular padding can incorporate all this information with a smaller convolution kernel (Figure 2c). We compared the receptive field of element *Y*_5,20_ under different convolution settings (Figure 2d), as well as the distribution of the coverage amount of each element’s receptive field under different convolution settings (Figure 2e). It can be seen that, compared to traditional convolution, diagonal convolution increases internal connections. Circular padding improves the issue of weak boundary receptive fields. And high dilation rates quickly enlarge the receptive field of each element, but also increase the non-uniformity of the receptive fields among different elements. Only the combination of diagonal convolution and high dilation rates can expand the receptive field while keeping it evenly distributed (Figure 2e).

To test the effects of high dilation rates and circular padding on diagonal convolution kernels, we experimented with various combinations of these elements: diagonal convolution kernels, high dilation rates, and circular padding. Specifically, we modified the type of convolution kernel, dilation strategy, and padding method within the same network structure, resulting in eight models: DiagConv, DiagConv + C, DiagConv + H, DiagConv + H + C, NaiveConv, NaiveConv + C, NaiveConv +H, NaiveConv + H + C, where “C” stands for circular padding, “H” for high dilation rate, “NaiveConv” for ordinary convolution kernels, and “DiagConv” for diagonal convolution kernels. It was observed that among all metrics, DiagConv + H + C achieved the optimal performance (Figure 2f). Comparing DiagConv and DiagConv + C revealed that adding circular padding alone did not significantly improve performance; however, a substantial enhancement was noticed when high dilation rates were also applied, which is related to the even distribution of the receptive field. On the other hand, comparing DiagConv with DiagConv + H indicated that without circular padding, using zero padding with high dilation rates led to a significant performance decline. A similar phenomenon occurred in all Naive models, though it was not as pronounced as in the DiagConv models. This is because ordinary convolution kernels lose a lot of intra-group information compared to diagonal convolution kernels. Therefore, when combined with diagonal convolution, neural networks are better able to leverage the advantages brought by high dilation rates and circular padding. Overall, our experiments demonstrated the effective synergy of diagonal convolution kernels, high dilation rates, and circular padding (Figure 2f).

We proposed a novel piecewise linear activation function, SoftSparse, which can yield sparse results (Figure 2g). The inspiration for SoftSparse comes from the soft thresholding operator (see Methods), and unlike the soft thresholding operator, SoftSparse does not lead to data scale collapse. We designed experiments to compare different activation functions, including SoftShrink (soft thresholding operator), SoftSparse, Tanh, ReLU, PReLU, and RReLU. Moreover, we introduced a new metric to measure flippability performance, the flip error (see Methods), and observed that all odd functions (SoftShrink, SoftSparse, Tanh) achieved the theoretical optimum value (0), whereas activation functions based on ReLU had higher flip errors (Figure 2g left). This validates the correctness of our flippability theory. At the same time, in terms of the AUC metric, SoftSparse and Tanh are top tier, while SoftShrink, due to causing data scale collapse, ranks slightly lower, and ReLU-based activation functions, due to their higher flip errors, perform the worst (Figure 2h right). This further validates our theory and the necessity of the design of the SoftSparse activation function.

### FSDiffNet enables efficient and fast inference in complex mixed distribution scenarios

We first conducted performance comparison experiments at a normal scale (*n* = 70, *p* = 39). Here, *n* is the number of samples of the observed data *X*, and *p* is the number of variables. We compared FSDiffNet with random matrices (Random), pseudo-inverse matrices (Pinv), and methods based on Gaussian likelihood such as GLasso^23^, GLasso*, JGL^24^, JGL*, Bayesian prior-based BDgraph^12^ and NetDiff^14^, along with two deep learning framework-based baseline models, Baseline_single and Baseline_double (Supplementary Materials). The asterisk (*) indicates the use of Gaussian likelihood for cross-validation parameter grid search tuning, which usually requires a significant amount of time. All single-condition methods (Pinv, GLasso, BDgraph, Baseline_single) calculated the graph matrices in their respective conditions to ultimately derive the differential network.

We randomly generated 100 differential graphs and used the observed data simulated from the latent graphs for inference. We employed the copula method to generate datasets with Gaussian distribution, exponential distribution, and mixed distribution (Figure S4). We calculated the AUPR, AUC, precision@1%, precision@5%, and time for these methods. In the comparison of performance in mixed distributions, we observed that FSDiffNet ranked first in both AUPR and Precision@5% (Figure 3a), and in the box plot of AUPR, it was noted that JGL, which relies heavily on Gaussian distribution, showed a significant decrease in performance after parameter tuning according to Gaussian likelihood (Figure 3b). We then aggregated all metrics (AUC, AUPR, Precision@1%, Precision@5%, −log(time)) and scaled each metric from 0 to 1 as a measure of relative performance across different methods, where 0 is the worst and 1 is the best method. The rankings were determined by average performance (Figure 3c). FSDiffNet achieved a score of 0.9, placing it first. In datasets with Gaussian and exponential distributions, the overall performance of FSDiffNet was comparable to the best-performing methods (Figures S11 and S12).

**Fig. 3:**
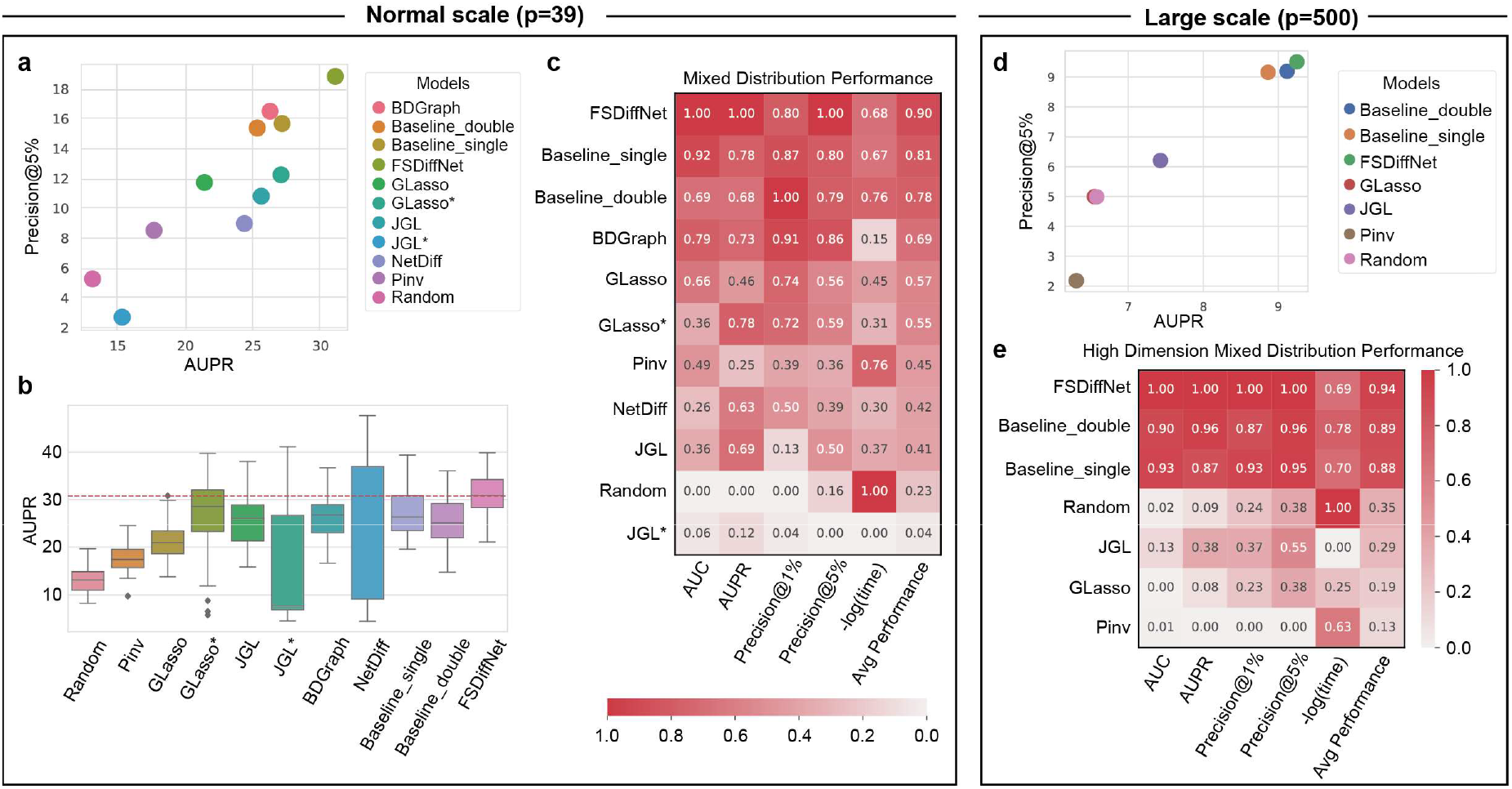
Comparison of performance among different methods. **a**. Performance comparison in a two-dimensional plot. The horizontal axis represents AUPR, and the vertical axis represents Precision@5%. **b**. Boxplot of AUPR for different methods (*n* = 100). **c**. Heatmap of comprehensive performance comparison under normal scale settings, with the horizontal axis representing different performance metrics and the last column representing average performance. The vertical axis lists the different methods, arranged in descending order of average performance, with the highest score being 1 and the lowest being 0. **d**. Two-dimensional performance comparison plot for *p* = 500, with the horizontal axis representing AUPR and the vertical axis representing Precision@5%. **e**. Heatmap of comprehensive performance comparison under large scale settings, similar to c.

We also compared the performance of different methods on challenging large-scale complex datasets, specifically under high-dimensional mixed distribution scenarios (*n* = 50, *p* = 500). Considering the time cost, we excluded comparisons with BDgraph, NetDiff, GLasso*, and JGL*. The results revealed that tasks in such scenarios are highly challenging; the straightforward method Pinv was completely ineffective, performing even worse than random matrices. The performance of GLasso was nearly equivalent to that of the random matrices Random (Figure 3d). It was observed that FSDiffNet, based on the deep learning framework, achieved significant performance improvements. Compared to JGL, FSDiffNet’s composite score reached 0.94, while JGL only scored 0.29.

We experimented with the time required by these methods to infer differential networks across datasets with different variable dimensions (Table 1). It was observed that the sampling-based method BDgraph requires a significant amount of time even for small values of *p*, and the time complexity of Bayesian inference-based methods increases sharply with an increase in *p*. However, when *p* increased to 2000 (requiring the inference of (*p*^2^ − *p*)/2 = 1,999,000 edges), FSDiffNet was still able to perform graph inference quickly and efficiently. Moreover, theoretical analysis can demonstrate that FSDiffNet is able to break through the computational complexity of *O*(*p*^3^), which is based on traditional statistical models and optimization algorithms (Supplementary Note 9).

**Table 1.** Comparison of the time complexity of different methods. In seconds (s).

| Methods | $p = 20$ | $p = 40$ | $p = 80$ | $p = 200$ | $p = 500$ | $p = 2000$ | time complexity |
| --- | --- | --- | --- | --- | --- | --- | --- |
| GLasso | 0.02 | 0.27 | 0.33 | 4.06 | 91.86 | 6847.58 | $O(p^3) \sim O(p^4)$ |
| JGL | 0.23 | 0.33 | 1.82 | 9.32 | 296.31 | — | $O(T_1 p^3)$ |
| BDgraph | 44.08 | 99.89 | 351.30 | 7592.43 | — | — | $O(T_2 p^5)$ |
| NetDiff | 4.84 | 36.4 | 289.49 | 4583.73 | — | — | $O(T_3 p^4)$ |
| FSDiffNet (CPU) | 0.08 | 0.10 | 0.32 | 0.99 | 4.27 | 56.32 | $O(p^2 \log(p))$ |
| FSDiffNet (GPU) | 0.03 | 0.04 | 0.04 | 0.07 | 0.28 | 4.53 | $O(p^2 \log(p))$ |

### FSDiffNet identifies the default mode network and its intrinsic connectivity changes in autism

In this experiment, we evaluated the effectiveness of FSDiffNet using functional Magnetic Resonance Imaging (fMRI) data from the Autism Brain Imaging Data Exchange (ABIDE) project^25^ and compared it with JGL and NetDiff. The ABIDE dataset provides a previously collected dataset of resting-state functional MRI data from 539 patients with autism spectrum disorder (ASD) and 573 control samples from all over the world. We used the nilearn Python package to download region of interest (ROI) annotations for the Multi-Subject Dictionary Study (MSDL) and datasets. Using the ABIDE data from Yale University, we obtained 22 autism group signal matrices and 19 healthy control group signal matrices, each with a shape of (169, 39), meaning 169 time points and 39 ROIs. We inferred differential functional connectivity maps for each pair of autism and control samples using different methods, resulting in a total of 418 predicted matrices, and calculated the average of these matrices as the final prediction of each method.

We first retained the top 10% of edges in the functional connectivity graph. Visually, the edges predicted by FSDiffNet are more concentrated and show stronger relationships. Predictions of other methods are more dispersed, exhibiting higher entropy (Figure 4a). Next, we calculated the regional strength by summing the absolute values of all neighboring edges for each node (Figure 4b). As described, the inferences of FSDiffNet show a more concentrated distribution of strengths, while other methods display a relatively flat distribution. Notably, the default mode network (DMN) was shown to be highly relevant to autism. According to the results, FSDiffNet successfully identified 3 out of 4 DMNs: the left DMN, right DMN, and medial DMN (Figure 4b bottom). The DMNs have been confirmed as key regions that undergo changes in individuals with autism^26–29^. Research has confirmed that in the ASD group, compared to the control group, activation of the Intraparietal Sulcus (IPS) during the motor sequence learning process is reduced^30^. Moreover, researchers have also found an abnormal increase in local connections between the occipital (Occ post) and temporal lobes in the ASD group^31^.

**Fig. 4:**
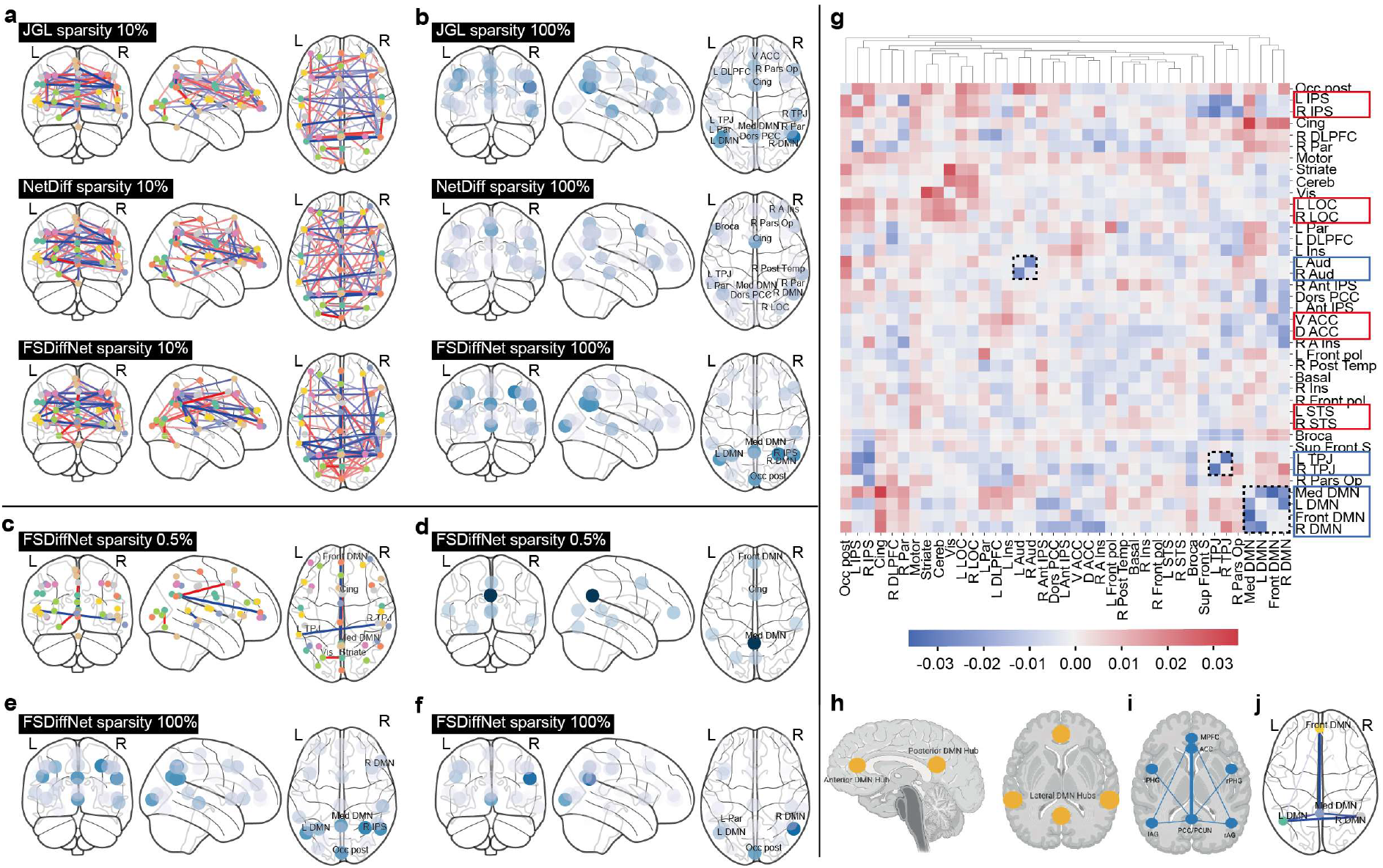
Differential brain functional connectivity graph inference results. **a**. The differential connectivity of the top 10% edges ranked by each method. **b**. The distribution of differential strengths for brain ROI inferred by each method, with strengths calculated from 100% of the differential edges. **c**. The differential connectivity of the top 0.5% edges ranked by FSDiffNet. **d**. The differential strength of brain ROI inferred by FSDiffNet, with strength calculated from the top 0.5% differential edges. **e**. The distribution of differential strengths for brain ROI inferred by FSDiffNet using the data from Yale University. **f**. The distribution of differential strengths for brain ROI inferred by FSDiffNet using the data from Stanford University. **g**. Heatmap of differential brain functional connectivity inferred by FSDiffNet, with the tree structure representing the results of hierarchical clustering. Red indicates connections that are enhanced in the ASD group, while blue indicates connections that are weakened in the ASD group. **h**. The four areas of the DMN. **i**. A review of the literature on patterns of weakened connections in ROI within ASD, with thicker lines indicating more literature support for the connection. **j**. The internal differential connectivity within the DMNs inferred by FSDiffNet, consistent with i.

Furthermore, we retained only the top 0.5% of edges, which corresponds to the 4 highest-weighted edges in the network (Figures 4c and d). The area with the largest difference is the medial DMN, which is connected to the anterior DMN and the Cingulate Cortex (Cing). The anterior cingulate cortex has been found to have reduced cognitive control over response inhibition in ASD, leading to repetitive behavior (RRB) in ASD ^32^. Posterior brain areas such as the primary visual cortex and the extra-striate cortex are more broadly activated in individuals with ASD, while frontal lobe areas are less active^33^. We also conducted the same experiment using the ABIDE data from Stanford University, which included 12 autism samples and 13 control samples. In Figure 4f (result of Stanford data), we can see similar results to Figure 4e (result of Yale data) described earlier, such as DMN and Occ post. This further confirms the robustness of FSDiffNet and the reliability of FSDiffNet’s discovery about ASD.

We performed hierarchical clustering on these areas using the differential connectivity features. In the inference results of FSDiffNet, we found that multiple regions with the same structural functions but different locations were clustered together, such as the left and right inferior parietal sulcus (L/R IPS), indicating that their differential patterns are similar (red box in Figure 4g). At the same time, we also found that in the group of autism patients, compared to the control group, some regions have weaker internal connections (indicated by the dashed boxes in Figure 4g), such as the left and right auditory cortices (L/R Aud). The reduction in connections between the auditory cortices (L/R Aud) in ASD patients may be related to their increased sensitivity to sounds. Moreover, the weakening of internal connections at the left and right temporoparietal junction (L/R TPJ) could be associated with social cognitive impairments in autism^34^. Notably, it can be observed that the internal connections within the DMN regions are significantly weakened in the ASD group. Some literature has verified that an underconnectivity pattern within intra-DMNs indeed occurs in the ASD group, especially between connections of the anterior and posterior DMN areas, with thicker lines representing more supporting literature (Figures 4h and i)^35^. This phenomenon is consistent with the results of FSDiffNet (Figures 4i and j).

### FSDiffNet identifies key driver genes differentiating breast cancer

We also compared FSDiffNet with other methods using the TCGA breast cancer dataset. In this analysis, we aimed to identify driver differential genes by inferring differential gene interaction graphs. Due to the large scale of the graph, we gave up the use of NetDiff for this part and instead adopted the Pearson Correlation Coefficient (PCC), a widely utilized and straightforward technique in the methods handling high-dimensional datasets. We downloaded the TCGA breast cancer RNA-seq gene expression data from the Xena data center, which was sequenced using the Illumina platform and normalized using FPKM. The dataset comprises 60,484 genes across 1,217 samples, including 1,104 cancerous and 113 normal samples. To reduce the dimensionality of the data, we identified the most relevant genes for breast cancer from KEGG database, selecting 1,228 candidate genes. Following quality screening, we narrowed this down to 500 genes exhibiting the highest variance, which we used for inferring the differential graph. This graph incorporates nearly 120,000 gene interactions, posing a significant challenge to most graph inference techniques.

To conduct a macro-level analysis, we initially preserved the top 500 edges of each inferred graph. Subsequently, we selected the main connected component from each method’s outputs as the definitive criterion for assessing the characteristics of the graphs. Intuitively, FSDiffNet’s results exhibited superior compactness and connectivity, whereas the JGL’s graph showed a more dispersed network distribution (Figure 5a). We calculated a range of topological characteristics for the series of graphs (Table 2). Owing to a substantial number of false positives, the graph generated by PCC was denser and more compact, comprising only 127 nodes. However, biological graphs are more likely to be scale-free graphs, which are sparse, heterogeneous, and moderately centralized. FSDiffNet has a relatively sparse graph structure (density of 0.028) and the most informative and centralized node distribution (heterogeneity of 1.418, centrality of 0.281). We also calculated the unit density centrality, i.e., the level of centrality under the same density condition, with FSDiffNet scoring 10.036. We attempted to fit the degree distribution of each method’s nodes to a power-law distribution. Figure 5b shows that FSDiffNet’s results are more consistent with a power-law distribution (coefficient of determination, *R*^2^ = 0.962), which is much higher than the other two methods.

**Table 2.** Comparison of the overall characteristics of the differential graphs. The most informative methods are marked in bold.

| Method | #nodes | #edges | D | R | CPL | Den. | Het. | Cen. | Cen./Den. |
| --- | --- | --- | --- | --- | --- | --- | --- | --- | --- |
| PCC | 127 | 499 | 8 | 4 | 3.082 | 0.062 | 1.090 | 0.267 | 4.306 |
| JGL | 361 | 467 | 26 | 13 | 8.290 | 0.007 | 0.844 | 0.068 | 9.714 |
| FSDiffNet | 187 | 492 | 7 | 4 | 3.223 | 0.028 | **1.418** | **0.281** | **10.036** |

D: Diameter, R: Radius, CPL: Characteristic path length, Den.: Density, Het.: Heterogeneity Cen.:Centralization, Cen./Den.:Centralization/Density.

**Fig. 5:**
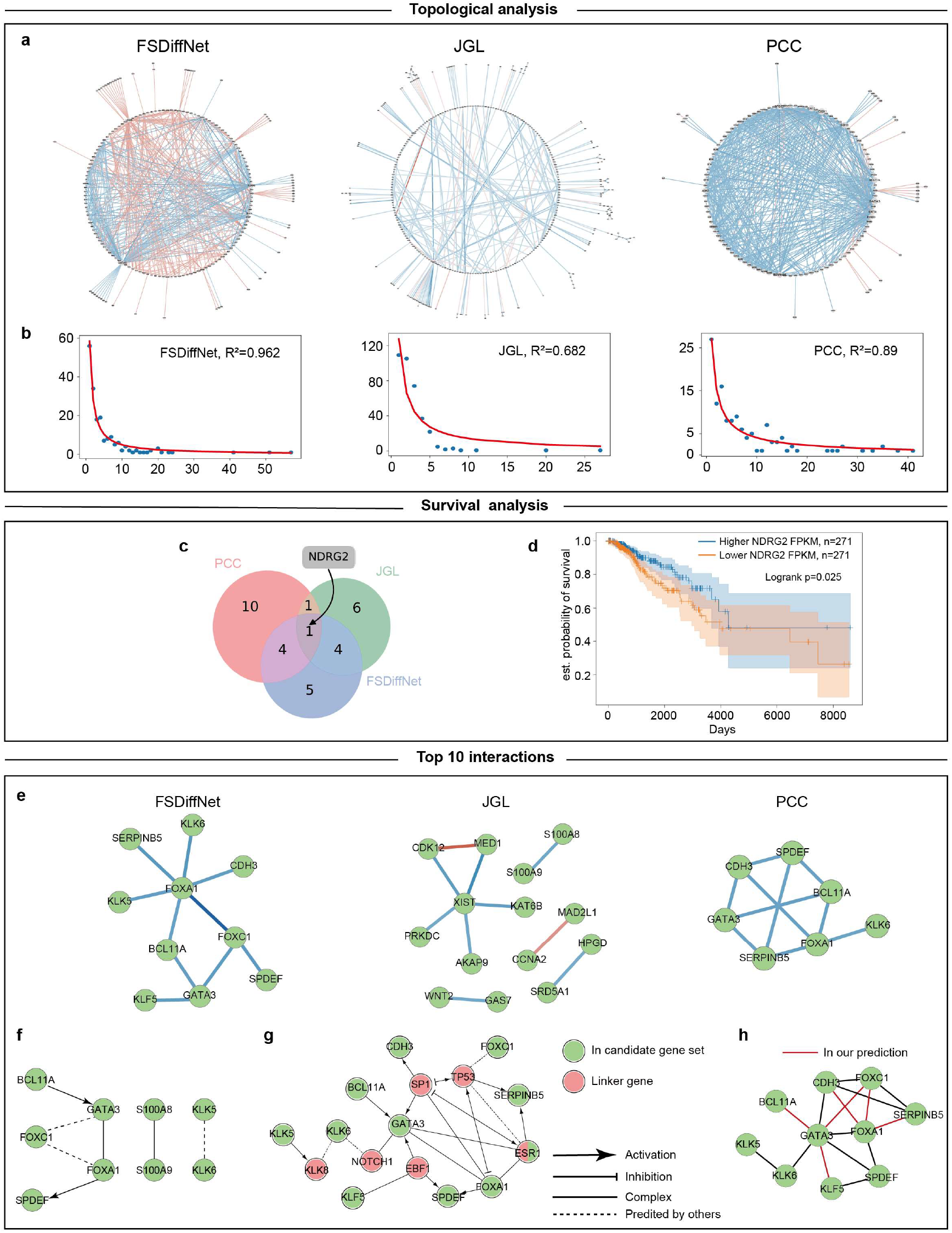
Breast cancer differential graph inference results. **a** Global structure of the differential gene interaction graph inferred by each method. The red edges indicate that the gene interaction strength in the cancer group is higher than that in the control group, and the blue edges indicate the opposite. **b** The power-law distribution fitting results. *R* is the the coefficient of determination. **c** Venn diagram of significant genes of various methods. **d** Survival analysis curve of NDRG2. The horizontal axis represents days, the vertical axis represents the estimated survival probability, and both the high-expression and low-expression groups include 271 samples each. **e** The subgraph composed of the top 10 edges ranked by each method. **f** Evidence interactions in reactome database. **g** Candidate gene interactions with linker genes in Reactome. The green nodes represent genes that are in our candidate gene set, and the red nodes represent linker genes, which did not appear in our input. **h** Contracted graph (merged red nodes in g).

Further, we selected the top 100 genes from each method as hub genes for survival analysis, which served as the gold standard to validate the effectiveness of these methods. The genes were ranked based on the sum of the absolute values of their edge weights. Specifically, we used the Kaplan-Meier model to fit survival curves and categorized the samples into “high expression” and “low expression” groups according to the gene expression levels. In this study, gene expressions above the 75th percentile were classified into the “high expression” group, while those below the 25th percentile were assigned to the “low expression” group. This approach allowed us to analyze the impact of genes with varying expression levels on patient survival times, thereby assessing the effectiveness of different methods in survival analysis. PCC identified 15 significant genes that potentially affect overall survival times. Meanwhile, JGL identified 12 key genes, and FSDiffNet found 14 significant genes (Table 3). We also calculated the overlapping genes between each pair of methods. The results show that FSDiffNet shared the most genes with either PCC or JGL (5 genes), while PCC and JGL only shared 2 genes (Figure 5c). This suggests that FSDiffNet, to some extent, integrates the advantages of both PCC and JGL. Among the findings, a common gene identified by all three methods was the NDRG2 gene. The survival analysis results for NDRG2 show that its log-rank p-value is 0.025, which is statistically significant (Figure 5d). NDRG2 is implicated as a potential tumor suppressor in breast cancer, primarily through its inhibitory effects on key oncogenic signaling pathways including Transforming Growth Factor-beta (TGF-β), Janus Kinase/Signal Transducer and Activator of Transcription (JAK/STAT), and Cyclooxygenase-2/Prostaglandin E2 (COX-2/PGE2). By modulating these pathways, NDRG2 significantly reduces the metastatic potential, cellular proliferation, and invasiveness of breast cancer cells, underscoring its therapeutic promise in mitigating breast cancer progression^36–41^. These conclusions are consistent with our survival analysis results, where high expression of NDRG2 was associated with enhanced survival probabilities for patients by inhibiting the progression of breast cancer.

**Table 3.** Comparison of the survival analysis results for the hub genes.

| Method (#genes) | Gene (log-rank p-values) |
| --- | --- |
| PCC (15) | CCL19 (0.001); CHI3L1 (0.014); CXCL2 (0.019); FABP7 (0.001); FGF1 (0.037); GREB1 (0.043); IGF1R (0.017); NDRG2 (0.025); PLK1 (0.042); RAC2 (0.002); RBBP8 (0.006); RELB (0.004); SEMA3B (0.000); SLC19A2 (0.028); WIF1 (0.013) |
| JGL (12) | CCL5 (0.030); CD69 (0.019); CXCL16 (0.014); JUND (0.017); KRT5 (0.007); LAPTM4B (0.039); NDRG2 (0.025); PDCD4 (0.017); RARRES1 (0.021); SDC1 (0.002); SLC19A2 (0.028); SLC5A8 (0.023) |
| FSDiffNet (14) | CHI3L1 (0.014); CXCL16 (0.014); FABP7 (0.001); FGF1 (0.037); KRT14 (0.023); KRT5 (0.007); LAPTM4B (0.039); LEF1 (0.004); NDRG1 (0.002); NDRG2 (0.025); RARRES1 (0.021); SFRP1 (0.050); WIF1 (0.013); WNT11 (0.032) |

To further explore gene interactions, we analyzed gene sets of the top 10 edges ranked by each method (Figure 5e). We treated the gene nodes covered by these edges as a list of genes, and verifiy potential associations using the Reactome database. We identified a total of seven gene interactions with high reliability in the database (Figure 5f). FSDiffNet successfully identified three established functional interactions; two of these are categorized as “regulation of expression” and involve the gene pairs BCL11A→GATA3 and FOXA1→SPDEF. An interaction involving the genes GATA3 and FOXC1 had been previously predicted by other research. In contrast, JGL identified only one existing “complex” interaction between the genes S100A8 and S100A9. Meanwhile, PCC only detected the interaction BCL11A→GATA3, which had also been identified by FSDiffNet.

To explore more detailed evidence, we utilized “linker” genes in the Cytoscape Reactome FI application. This approach introduces genes that are not present in the candidate gene set as first-order neighbors, allowing for the creation of a more extensive potential linkage map (Figure 5g). To more visually display the results driven by data, we constructed a contracted graph. Specifically, if the shortest path between two candidate genes (marked in green) included only linker genes, we connected these two nodes, thereby forming a contracted graph (Figure 5g). The results show that 6 out of the 10 edges identified by FSDiffNet (60%) are present in the contracted graph, with FOXA1 and GATA3 still appearing as central nodes in the differential graph, implying they may primarily mediate the development of breast cancer. Research indicates that FOXA1 mutations are considered a hallmark of ER+ breast cancer and may affect treatment responses ^42^. GATA3 has also been proven to be an independent prognostic marker for breast cancer, with its low expression levels always associated with poorer prognosis and cancer recurrence ^43^.

## Discussion

This paper primarily introduces FSDiffNet, a flippable siamese neural network based on a deep learning framework. We first proposed the concept of flippability, analyzed the theoretical properties of flippable mappings, and delineated the necessary conditions for constructing a flippable neural network (Theorem 2). We designed the SoftSparse activation function for nonlinear sparse mapping and presented the practical architecture of FSDiffNet. On challenging mixed distribution datasets, FSDiffNet can surpass the performance of existing methods in a very short time. The theoretical analysis of time complexity demonstrates that FSDiffNet can break through the computational complexity of traditional optimization model algorithms *O*(*p*^3^), and experiments verified that FSDiffNet can utilize GPUs to complete differential graph inference for 2000-dimensional variables within 5 seconds. Finally, in experiments on datasets of patients with autism, FSDiffNet successfully recovered the differential functional connectivity maps of ROIs, particularly identifying weakened connection patterns in the DMNs within the ASD group. In the breast cancer dataset, FSDiffNet inferred large-scale graphs and successfully identified the NDRG2 gene closely related to survival analysis, aiding researchers in better understanding the driving differences in disease.

## Methods

### Problem Definition

Given two sets of observation data matrices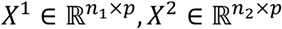, generated from the underlying undirected graph structures *G*^1^ = (*V, E*^1^, *W*^1^) and *G* = (*V, E*^2^, *W*^2^) respectively, where *V* is the set of vertices (variables), *E*^1^, *E*^2^ are the sets of edges, and *W*^1^, *W*^2^ are the edge weight matrices. Here, *V* is known and shared between the two conditions, whereas *E*^1^, *W*^1^, *E*^2^, *W*^2^ are unknown. The number of vertices is *p* = |*V*| and the number of observation samples are *n*_1_, *n*_2_ for the two conditions respectively. 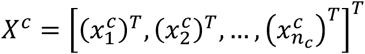 where 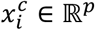 is the *i*-th observation sample from the *c*-th group, represented as a *p*-dimensional vector. The differential graph inference problem is defined as finding a mapping 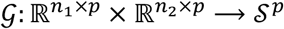,where 𝒮^p^ is the space of *p*-dimensional real symmetric matrices, such that 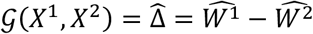 is as close as possible to the true 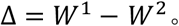

### Empirical Risk Function

Our goal is to construct a 𝒢 as described in the problem definition based on neural networks. We can consider the problem as a classification task:

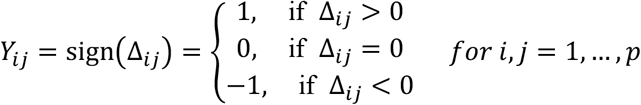

Here, *Y*_*ij*_ represents the classification label for the edge (*i, j*). *Y*_*ij*_ = 1 indicates a positive effect, *Y*_*ij*_ = −1 indicates a negative effect, and if there is no edge, then *Y*_*ij*_ = 0. Unlike ordinary classification problems, as a graph inference algorithm, we classify *p* × *p* edges simultaneously. This is because these edges are interdependent, unlike in general classification problems where classification targets are independent. The empirical risk function is defined as follows:

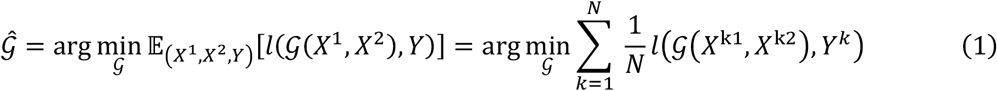

where *l*: ℝ^*p*×*p*^ × ℝ^*p*×*p*^ → ℝ is a loss function designed by the user based on the actual problem context. It is worth noting that some traditional methods can be viewed as selecting 𝒢 within a constrained mapping space, such as 𝒢 (*X*^1^, *X*^2^) = *corr*_*pcc*_(*X*^1^) − *corr*_*pcc*_ (*X*^2^), where *corr*_*pcc*_ corresponds to methods based on the Pearson correlation coefficient.

### Extended Loss Function

For multi-classification problems, a categorical cross-entropy loss function is commonly used. However, since our labels contain coupled information, where the positive and negative effects are in a dual relationship, we extend the binary cross-entropy loss function to obtain the following loss function:

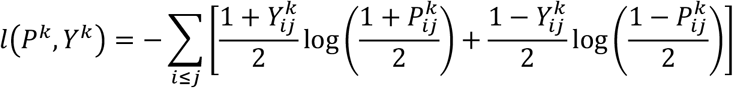

where *P*^k^ = 𝒢 (*X*^k1^, *X*^k2^). Especially, due to the sparsity of the graph leading to label imbalance in this task, we introduce a focal factor *α* > 0 with the aim of making the edges’ positions easier to learn. The total loss function formula is as follows:

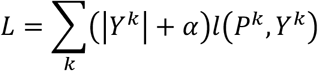

### Flippability and Its Theory

#### Definition 1 (Flippability)

For a mapping 𝒢: 𝒳 × 𝒳 → 𝒮 over a totally ordered space 𝒳, and ∀*X*^1^, *X*^2^ ∈ 𝒳, if it satisfies 𝒢(*X, X*) = −𝒢(*X, X*), then we say 𝒢 is flippable or 𝒢 possesses flippability. Through the definition of flippability, we present the properties of a flippable mapping:

##### Theorem 1

Let 𝒳 be a totally ordered space. Given any flippable mapping 𝒢: 𝒳 × 𝒳 → 𝒮, there exists a mapping *f*: 𝒳 × 𝒳 → 𝒮 such that 𝒢(*X*^1^, *X*^2^) = *f*(*X*^1^, *X*^2^) − *f*(*X*^2^, *X*^1^).

##### Property 1

Given a flippable mapping 𝒢: 𝒳 × 𝒳 → 𝒮, and a sequence of mappings *g, g*, … *g*_k_: 𝒮 → 𝒮. Then *g*_1_ ∘ *g* _2_∘ ⋯ ∘ *g*_k_ ∘ 𝒢 is flippable if and only if *g*_1_, *g*_2_, …, *g*_k_ are all odd mappings.

##### Theorem 2

Assume ℱ is a flippable neural network, then there must exist a prime flippable network ℱ_O_ such that ℱ = *g*_1_ ∘ *g*_2_ ∘ ⋯ ∘ *g*_k_ ∘ ℱ_O_, where *g*_1_, *g*_2_, …, *g*_k_ are all odd network layers (odd mappings).

The detailed explanations and proofs of the above theorems are presented in Supplementary Materials.

### Flippability theory guides component and architecture design for FSDiffNet

Addressing the flip problem in differential graph inference using neural network architectures is a significant challenge. This problem demands that the neural network 𝒢, satisfies the condition 𝒢(*X*^1^, *X*^2^) = −𝒢(*X*^2^, *X*^1^). Achieving such flippability can be easily realized in manually designed 𝒢, such as in sparse optimization inference methods based on Gaussian graphical models. However, obtaining this flippability within the framework of neural networks is not straightforward. A trivial approach involves training a single-condition inferencer *f*, and then subtracting the results for two conditions, that is 𝒢(*X*^1^, *X*^2^) = *f*(*X*^1^) − *f*(*X*^2^). However, this method loses the coupled information between the two conditions, rendering joint inference challenging. If one aims to input data from both conditions for joint learning and inference, the data typically loses its flippability after traversing several neural network layers, resulting in outputs that often lack information regarding the original order of the input channels.

To overcome this limitation in neural networks, we have proposed the definition of flippability and elucidated the properties of flippable mappings, thereby establishing the necessary theoritical conditions for constructing a flippable neural network within the framework of neural networks. These theories have aided us in designing the network architecture. Theorem 1 elucidates the decomposability of flippable mappings, while Property 1 clarifies the extension conditions for flippable mappings. Consequently, by restricting the mapping within the structure of a neural network, we present an important theorem, Theorem 2, which assists in designing a flippable neural network architecture.

Theorem 2 suggests that to construct a flippable neural network ℱ, it is necessary to design intermediate odd neural network layers, {*g*_*l*_}_*l*=1,…_, such as odd convolutional layers, activation function layers, or batch normalization layers, as well as a flippable subnetwork ℱ_0_. Theorem 1 informs us that for any flippable mapping 𝒢, there exists at least one *f* such that 𝒢(*X*^1^, *X*^2^) = *f*(*X*^1^, *X*^2^) − *f*(*X*^2^, *X*^1^), which is the form of a siamese network. With these theoretical results, we can naturally design and achieve the neural network architecture of FSDiffNet.

### Diagonal Convolution

Building upon DeepGraph^15^, we introduced the concept of Diagonal Convolution kernels (DiagConv, DC). Specifically, traditional convolution kernels can be represented as follows:

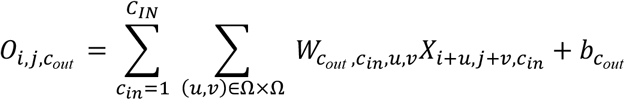

where 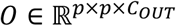 is the output of the convolution layer, 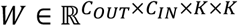 is the weight matrix, 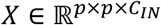 is the input tensor, 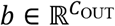 is the bias term, *C*_*IN*_, *C*_*OUT*_ are the numbers of input and output channels, respectively, and *K* is the size of the convolution kernel, set to 3 in this work. 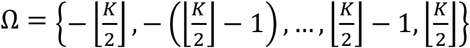 the set of weight indices.

Similarly, we define diagonal convolution in the following form:

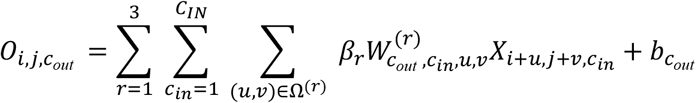

where Ω^(1)^ = (*d*_*l*_Ω) × (*d*_*l*_Ω), Ω^(2)^ = (*j* − *i* + *d*_*l*_Ω) × (*d*_*l*_Ω), Ω^(3)^ = (*d*_*l*_Ω) × (*i* − *j* + *d*_*l*_Ω) are the sets of weight indices, *d*_*l*_ is the dilation rate of the *l*−th layer network, and *β*_*r*_ is the convolution kernel weight for the *r*-th part. Compared to traditional convolution kernels, DiagConv covers not only external connections but also includes nodes’ internal connections. In this paper, each sub-part is shaped as a 3*3 kernel, totaling 27 weights in one channel.

### High Dilation Rate

A method to rapidly cover all elements in a matrix on another scale with fewer layers, while keeping the kernel size fixed, is to increase the dilation rate of the convolutional kernel^44^. For traditional convolutional kernels, the dilation rate is 1. In this paper, we set the dilation rate as a sequence related to the layer number *l*, {3^*l*^}_*l*=O,…,*L*_. Through this setting, the convolutional kernel can cover all elements as quickly as possible without missing any elements (Supplementary Materials).

### Circular Padding

The primary challenge with setting a high dilation rate is that the convolutional kernel can easily exceed image boundaries, which necessitates an appropriate padding technique to adjust the convolution. Zero padding is the most commonly used method^15^. However, zero padding does not suit this kind of problem well. We begin with *Y*, where the classification scores *Ŷ* predicted by deep learning frameworks are typically in decimal form, with their absolute values indicating prediction probabilities. Hence, we can link them to the normalized matrix *P* of the precision matrix *Θ*, defined as 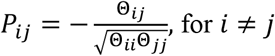.

Noting that the precision matrix 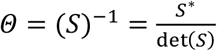 is the inverse of the covariance matrix *S*, where *S*^*^ is the adjugate matrix of *S* ^45^. Thus, whether *Θ*_*ij*_(or *Y*_*ij*_)is zero is dependent on 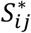 (for instance, 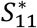 calculated using *S* _22_, *S* _23_, *S* _32_, *S*_33_ in Figure 2b). If the objective is to cover all necessary elements with fewer layers, circular padding is the most effective technique.

### SoftSparse Activation Function

Based on Theorem 2, we need to design odd activation function layers. Hence, we introduced a novel piecewise linear activation function, SoftSparse, which can produce sparse results. The inspiration for SoftSparse comes from solving the subproblems of the joint graphical Lasso model:

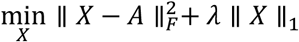

The closed form of this problem can be obtained by the soft-thresholding function^46^:

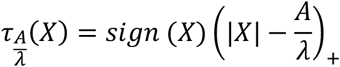

where (·)_+_ is the positive part function, and *sign* is the sign function. The intuitive understanding is that this function maps *X*_*ij*_ to 0 if I*X*_*ij*_I is less than the threshold; otherwise, it subtracts the threshold from the magnitude of *X*_*ij*_. This causes the image space of *τ* to be a reduction from its domain space, i.e., *Im*(*τ*) ⊂ *Dom*(*τ*). Even though *τ* is an odd mapping, using *τ* as an activation function would progressively shrink the data scale with the advancement of network layers, possibly even leading to data disappearance. Therefore, we designed SoftSparse, *σ*_*λ*_ (*X*) = *X*𝕀_> *λ*_ (|*X*|), where 𝕀 is the indicator function. This activation function is not only nonlinear (piecewise linear) but also maintains data scale, achieving sparse results. SoftSparse smoothens those elements whose absolute values are less than the threshold *λ* while ReLU smoothens elements less than 0, as shown in Figure 2g.

### Processing in Batch Normalization

Batch normalization^47^ is a technique used to standardize the inputs to a neural network layer, which can speed up training and improve model performance. It is typically applied before the activation function layer. Therefore, in addition to the odd activation function layers, the batch normalization layer also needs to be addressed. The general form of a batch normalization layer is as follows:

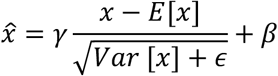

where 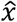 is the output, *x* is the input, *γ, β* are learnable affine parameters that allow for an affine transformation of the batch-corrected output. If momentum is incorporated during training for tracking statistics, the form is as follows:

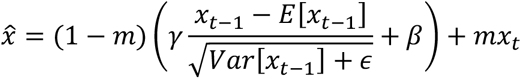

where *m* is the momentum term parameter, typically set to 0.1, *x*_*t*_ is the data observed in the new batch, and *x* _*t* − 1_is the data observed in the previous batch. Therefore, if the batch normalization layer is required to be an odd mapping, it is necessary to set *β* = 0. This can be achieved by setting affine=False in PyTorch. The processed batch normalization layer form is as follows:

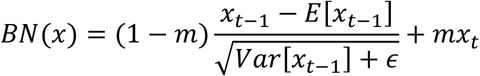

### Representation of FSDiffNet

The FSDiffNet is not a mere sequential network structure; it comprises three parallel channels. Here, we use *f* to denote one of the channels in the siamese network, and ℱ to represent the channel at the bottom (as shown in Figure 2a). The inputs to FSDiffNet are two correlation coefficient matrices *S*^1^, *S*^2^. The zeroth layer of ℱ is given by:

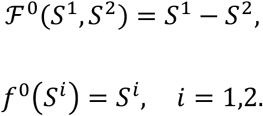

The first layer of ℱ is given by:

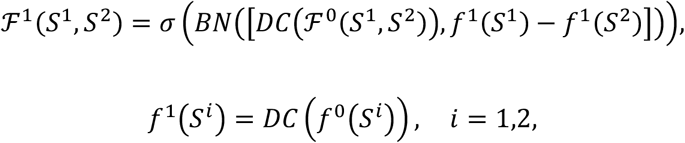

where *σ* represents the SoftSparse activation layer, *DC* is the bias-free diagonal convolution,BN is odd batch normalization layer, and [·] denotes the concatenation operation.

The *l*-th layer of ℱ can be expressed as:

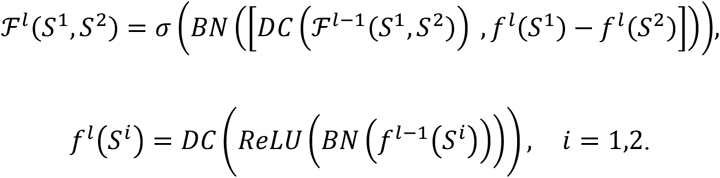

The second-to-last layer of ℱ can be expressed as:

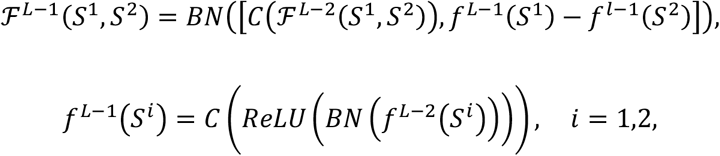

where *C* is the ordinary bias-free convolution layer with 1*1 kernel. The final output layer of ℱ can be expressed as:

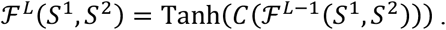

### Evaluation Metrics

In this study, we employed four key performance evaluation metrics: AUC, AUPR, Precision@1%, and Precision@5%. AUC, or the area under the ROC curve, measures the overall performance of a model across different classification thresholds, with values closer to 1 indicating better model performance. AUPR, or the area under the precision-recall curve, is suited for evaluating model performance in imbalanced datasets, especially when the number of positive samples is significantly less than that of negative samples, as it can more accurately reflect the model’s actual performance. Precision@1% focuses on the model’s prediction accuracy for the top 1% of data that are most likely to be positive samples, while Precision@5% evaluates the accuracy for the top 5% of data, based on the same ranking criteria. Both are used to assess the model’s ability to identify highly probable positive samples in extremely imbalanced data environments. Particularly, since the labels involve both positive and negative correlations, it is necessary to separately evaluate the metrics for each and then calculate the average.

### Flip Error

To measure the degree of flippability violation, we propose the definition of flip error as follows. Given a sequence of 2N test samples, 𝕊 = {(*X*^*i*, 1^, *X*^*i*, 2^), (*X*^*i*, 2^, *X*^*i*, 1^)}_*i*=1,…,*N*_, and an estimator ℱ: 𝒳 × 𝒳 → 𝒮, the flip error can be written in the following form:

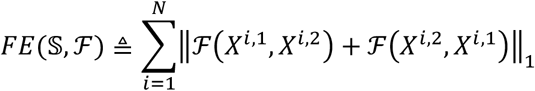

It can be observed that the theoretical lower bound of the flip error is 0. The neural network we designed is capable of strictly achieving this lower bound (see Figure 2).

### Statistical Information

In this study, we utilized the log-rank test^48^ to assess the differences in survival times between two independent sample groups (see Figure 5), where the key statistical indicator is the log-rank p-value. The log-rank test is a non-parametric method that evaluates whether survival curves are significantly different by comparing the differences in survival rates at each observed time point across groups. Specifically, the statistic for the log-rank test can be calculated using the following formula:

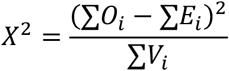

Here, *O*_*i*_ is the number of events (such as deaths or recurrences) observed at the *i* th time point, *E*_*i*_ is the expected number of events at the *i* th time point under the assumption that the two groups have the same survival function, and *V*_*i*_ is the variance at the *i* th time point. The calculated *X*^2^ value is used to derive the log-rank p-value. If the p-value is less than a predetermined level of significance (usually 0.05), it indicates a statistically significant difference in survival times between the two groups, implying that treatment effects or survival statuses vary across different groups.

## Data Availability

All data related to brain graph reconstruction in this paper can be downloaded through the ‘datasets’ module in the ‘nilearn’ Python package. For specific tutorials, the readers can refer to: https://nilearn.github.io/. All breast cancer-related data used in this paper can be downloaded from the UCSC Xena^49^ database by using the keyword ‘TCGA-BRCA’ at the link: https://xenabrowser.net/datapages/.

## Code Availability

To facilitate use by researchers, we have packaged the code into a Python package named ‘fsdiffnet’, which can be easily accessed by consulting the documentation at https://github.com/amssljc/FSDiffNet.

## Ethics declarations

### Competing interests

The authors declare no competing interests.

## Supplementary Information

See Supplementary Materials.

## Notes

### Competing Interest Statement

The authors have declared no competing interest.

