## Supplementary Materials for "Flippable Siamese Differential Neural Network for Differential Graph Inference"

### 1. Supplementary Note 1: Details of Related Works

#### 1.1 Background knowledge

##### 1.1.1 MLE-based sparsity optimization

Probability graphical models, especially Gaussian graphical models were used to infer underlying graph structure for a long while. One of the single-condition classical models is Gaussian Graphical Lasso [1,2], which estimates the precision matrix $\widehat{\Theta}$ as follows:

$$\widehat{\Theta} = \arg \min_{\Theta \in \mathcal{S}_{++}^{p}} -n \log \det \Theta + n\, tr(S\Theta) + \lambda \|\Theta\|_1$$

where $n$ is the sample number, $S$ is the empirical sample covariance matrix, $\lambda$ is the tuning parameter to control the sparsity of the graph adjacency matrix $\Theta$, $\|\cdot\|_1$ is the $L1$ norm which is the sum of the absolute values of the matrix elements, $tr(\cdot)$ is the trace of a matrix, and $\mathcal{S}_{++}^{p}$ represents the symmetric positive definite matrix space. $-L(\Theta, S) = -n \log \det \Theta + n\, tr(S\Theta)$ is the negative likelihood function and $P(\Theta) = \lambda\|\Theta\|_1$ is the sparsity penalty. The main idea of this model is to balance $L(\Theta)$ with $P(\Theta)$. The more likelihood information used, the less the sparsity is. The most straightforward way to obtain the differential graph is to infer the two graphs of different conditions and obtain $\widehat{\Delta} = \widehat{\Theta^1} - \widehat{\Theta^2}$ as the result. However, this method loss much of information in data. The more rational way to infer $\Delta$ is to joint estimate $\Theta^1, \Theta^2$ with a joint penalty as follows [3–10]:

$$\widehat{\Theta^1}, \widehat{\Theta^2} = \arg \min_{\Theta^1, \Theta^2 \in \mathcal{S}_{++}^{p}} \sum_{i=1}^{2} \left( -L(\Theta^i, S^i) + \lambda_1 \left\|\Theta^i\right\|_1 \right) + \lambda_2 P_2(\Theta^1, \Theta^2)$$

where $P_2(\Theta^1, \Theta^2)$ is the joint penalty term that usually controls the sparsity of the differential graph. For example, $P_2(\Theta^1, \Theta^2) = \|\Theta^1 - \Theta^2\|_1$ in the fused graphical lasso (FGL) model, and $P_2(\Theta^1, \Theta^2) = \sum_{i \neq j} \left\| \begin{bmatrix} \Theta_{ij}^1 \\ \Theta_{ij}^2 \end{bmatrix} \right\|_2$ in the group graphical lasso (GGL) model [11]. Another approach is to direct estimate $\Delta$ as a variable which can be described as follows [12,13]:

$$\widehat{\Delta} = \min_{\Delta} L_p(\Delta) + \lambda_1 \|\Delta\|_1$$

where $L_p(\Delta)$ is the pseudo-likelihood function, and $\lambda_1$ controls the sparsity of the differential graph. $L_p(\Delta)$ is designed as a convex function with gradient $\nabla L_p(\Delta) = (S^1 - S^2)^{-1} - \Delta = 0$. For example, $L_p(\Delta)$ is designed to be $\frac{1}{2} tr(\Delta S^1 \Delta S^2) - tr\left(\Delta(S^1 - S^2)\right)$[14]. But MLE-based methods usually more depend on the likelihood function, and its background assumption, such as multivariate-Gaussian distribution in the GLasso model.

1.1.2 Bayes-based MCMC sampling inference

Another kind of approach to infer differential graph maximizes the posterior probability of $\Theta$, rather than directly maximize s the likelihood function [15–17]. Generally, the joint posterior distribution can be written as follows:

$$\mathbb{P}(G^1, G^2, \Theta^1, \Theta^2 | X^1, X^2) \propto \mathbb{P}(X^1, X^2 | G^1, G^2, \Theta^1, \Theta^2)\mathbb{P}(\Theta^1, \Theta^2 | G^1, G^2)\mathbb{P}(G^1, G^2)$$

where $\mathbb{P}(G^1, G^2)$ is usually a uniform distribution, $\mathbb{P}(X^1, X^2 | G^1, G^2, \Theta^1, \Theta^2)$ is the likelihood function same as that in section 1.1. $\mathbb{P}(\Theta^1, \Theta^2 | G^1, G^2)$ is usually a G-Wishart distribution [18,19] as follows:

$$\mathbb{P}(\Theta^1, \Theta^2 | G^1, G^2) = \frac{1}{I(b, B)} \prod_{i=1}^{2} \left(\Theta^i\right)^{\frac{b-2}{2}} \exp\left\{-\frac{1}{2} tr\left(\Theta^i B\right)\right\}$$

where $b > 2$ is the degree of freedom, and $B$ is a positive symmetric definite parameter matrix. $I(b, B) = \int_{\mathcal{G}^1} \int_{\mathcal{G}^2} \prod_{i=1}^{2} \left(\Theta^i\right)^{\frac{b-2}{2}} \exp\left\{-\frac{1}{2} tr\left(\Theta^i B\right)\right\} d\Theta^1 d\Theta^2$ is the normalizing constant, $\mathcal{G}^i$ is the graph space of group $i$. BDgraph considers the graph inference problem as a Birth-Death (BD) model by updating the posterior probabilities and proposes a BDMCMC method [16]. NetDiff calculate the Bayes factor $\beta = \frac{\mathbb{P}(X^2 | G^2)}{\mathbb{P}(X^1 | G^1)}$, where $\mathbb{P}\left(X^i | G^i\right) = \int_{\mathcal{G}^i} \mathbb{P}\left(X^i | G^i, \Theta^i\right)\mathbb{P}\left(\Theta^i | G^i\right) d\Theta^i$ is calculated by MCMC sampling method [17]. However, the main weakness of this kind of methods is that they are depending on the prior distributions and the huge amount of time cost.

#### 1.2 Implementation details of the comparison methods

For the ‘Random’ method, we utilized ‘torch.rand’ function generates random matrices with outputs constrained between -1 and 1. For the ‘Pinv’ method, we used the ‘np.linalg.pinv’ function to calculate the pseudo-inverse matrix of the correlation coefficient matrix provided for each condition, serving as the predicted precision matrices $\widehat{\Theta^1}, \widehat{\Theta^2}$. For the GLasso method, we employed the ‘GraphicalLasso’ class from Python module ‘sklearn.covariance’, with default parameters (alpha=0.01). GLasso* utilized the ‘GraphicalLassoCV’ class from ‘sklearn.covariance’ for automatic grid cross-validation. For the JGL method, due to the lack of a Python implementation, we used the ‘rpy2’ Python package to call the R package ‘JGL’ within Python, adopting default parameters (lambda1=0.1, lambda2=0.1, penalty="fused"). For JGL*, we implemented a cross-validation functionality similar to ‘GraphicalLassoCV’ and named it ‘JointGraphicalLassoCV’. Similarly, for the BDGraph and Netdiff methods, we used rpy2 to call the R packages ‘BDgraph’ and ‘NetDiff’, respectively, using default parameters according to official documentation. All these methods can be accessed by installing our published Python package ‘difflearn’, available at: https://github.com/amssljc/difflearn.

### 2. Supplementary Note 2: Flippability Theory

#### 2.1 Flippability Definition

**Definition 1 (Flippability).**

For a mapping $\mathcal{G}: \mathcal{X} \times \mathcal{X} \to \mathcal{S}$ over a totally ordered space $\mathcal{X}$, and $\forall X^1, X^2 \in \mathcal{X}$ , if it satisfies $\mathcal{G}(X^1, X^2) = -\mathcal{G}(X^2, X^1)$ , then we say $\mathcal{G}$ is flippable or $\mathcal{G}$ possesses flippability.

Here, we abstract $\mathcal{X}, \mathcal{S}$ from the sample space to a more general space to obtain more universal results. Unless specified, they can refer to any algebraic space with linear mapping. Due to symmetry, without loss of generality, we only consider the case where $X^1$ is in front.

**Corollary 1** If $\mathcal{G}$ is flippable, then for all $X^1 \in \mathcal{X}$, $\mathcal{G}(X^1, X^1) = 0$. Where 0 is the zero element in $\mathcal{S}$.

**Lemma 1** For any given mapping, $f: \mathcal{X} \times \mathcal{X} \to \mathcal{S}$, having $\mathcal{G}(X^1, X^2) = f(X^1, X^2) - f(X^2, X^1)$ is flippable. Specifically, for any given mapping $f: \mathcal{X} \to \mathcal{S}$, having $\mathcal{G}(X^1, X^2) = f(X^1) - f(X^2)$ is flippable.

**Definition 2** (**Flippable Partition**). Given a space $\mathcal{X} \times \mathcal{X}$ and two sets within the space $\Gamma_a, \Gamma_b \subset \mathcal{X} \times \mathcal{X}$, if $\Gamma_a, \Gamma_b$ satisfy the following conditions:

a) $\Gamma_a \cup \Gamma_b = \mathcal{X} \times \mathcal{X}$;

b) For all $(X^1, X^2) \in \Gamma_a$, it holds that $(X^2, X^1) \in \Gamma_b$;

c) For all $(X^1, X^2) \in \Gamma_b$, it holds that $(X^2, X^1) \in \Gamma_a$;

d) For all $(X^1, X^2) \in \Gamma_a \cap \Gamma_b$, it holds that $X^1 = X^2$;

Then $\{\Gamma_a, \Gamma_b\}$ is called a flippable partition of $\mathcal{X} \times \mathcal{X}$. For example, let $\mathcal{X} = \mathbb{R}$, $\Gamma_a = \{(x, y) | y \geq x \,\&\, x, y \in \mathbb{R}\}$, $\Gamma_b = \{(x, y) | y \leq x \,\&\, x, y \in \mathbb{R}\}$, $\Gamma_a \cap \Gamma_b = \{(x, y) | y = x \,\&\, x, y \in \mathbb{R}\}$.

**Definition 3 (Total Order Relation).**

For a space $\mathcal{X}$, for all $\forall a, b, c \in \mathcal{X}$, if a binary relation $\leq$ satisfies the following conditions:

- Transitivity: If $a \leq b, b \leq c$, then $a \leq c$;
- Completeness: Either $a \leq b$ or $b \leq a$;
- Antisymmetry: If $a \leq b$ and $b \leq a$, then $a = b$;

Then the space $\mathcal{X}$ is said to satisfy the total order relation $\leq$.

**Definition 4 (Total Order Space).**

A space that satisfies the total order relation is called a total order space.

**Corollary 2** For any total order space $\mathcal{X}$, there exists a flippable partition of $\mathcal{X} \times \mathcal{X}$.

**Proof:** Let $\mathcal{X}$ satisfy the total order relation $\leq$. Define $\Gamma_a = \{(X^1, X^2) | X^2 \leq X^1 \ \& \ X^1, X^2 \in \mathcal{X}\}$, $\Gamma_b = \{(X^1, X^2) | X^1 \leq X^2 \ \& \ X^1, X^2 \in \mathcal{X}\}$. By the completeness of the order, part a) is proven. Parts b) and c) are naturally proven by the definition of $\Gamma_a, \Gamma_b$. By the antisymmetry of the order, part d) is proven. Therefore, $\{\Gamma_a, \Gamma_b\}$ is a flippable partition of $\mathcal{X} \times \mathcal{X}$. ✅

#### 2.2 Properties of Flippable Mappings

**Theorem 1** Given $\mathcal{X}$ as a total order space, for any flippable mapping $\mathcal{G}: \mathcal{X} \times \mathcal{X} \to \mathcal{S}$, there exists a mapping $f: \mathcal{X} \times \mathcal{X} \to \mathcal{S}$ such that $\mathcal{G}(X^1, X^2) = f(X^1, X^2) - f(X^2, X^1)$.

**Proof:** Let

$$f(X^1, X^2) = \begin{cases} \mathcal{G}(X^1, X^2) & , (X^1, X^2) \in \Gamma_a \\ 0 & , (X^1, X^2) \notin \Gamma_a \end{cases}$$

where $\{\Gamma_a, \Gamma_b\}$ is a flippable partition of $\mathcal{X} \times \mathcal{X}$. According to Corollary 1 and Definition 2, it can be concluded that $\mathcal{G}(X^1, X^2) = f(X^1, X^2) - f(X^2, X^1)$. ✅

**Lemma 2** Given a flippable mapping $\mathcal{G}: \mathcal{X} \times \mathcal{X} \to \mathcal{S}$ and a mapping $g: \mathcal{S} \to \mathcal{S}$, then $g \circ \mathcal{G}$ is flippable if and only if $g$ is an odd mapping.

**Proof:** For all $X^1, X^2 \in \mathcal{X}$, $g \circ \mathcal{G}(X^1, X^2) = g\big(\mathcal{G}(X^1, X^2)\big) = g\big(-\mathcal{G}(X^2, X^1)\big) = -g\big(\mathcal{G}(X^2, X^1)\big) = -g \circ \mathcal{G}(X^2, X^1)$ if and only if $g$ is odd. ✅

**Corollary 3** Given an odd mapping $g: \mathcal{X} \to \mathcal{S}$, $\mathcal{G}(X^1, X^2) = g(X^1 - X^2)$ is flippable.

**Corollary 4** The concatenation of two flippable mappings $\mathcal{G}_1, \mathcal{G}_2$ into $\mathcal{G}(X^1, X^2) = [\mathcal{G}_1(X^1, X^2), \mathcal{G}_2(X^1, X^2)]$ remains flippable.

**Corollary 5** Given a flippable mapping $\mathcal{G}: \mathcal{X} \times \mathcal{X} \to \mathcal{S}$ and a sequence of mappings $g_1, g_2, \dots g_k: \mathcal{S} \to \mathcal{S}$. Then $g_1 \circ g_2 \circ \cdots \circ g_k \circ \mathcal{G}$ is flippable if and only if each of $g_1, g_2, \dots, g_k$ is an odd mapping.

##### 2.3 Prime-Flippable Networks

This section continues the discussion on neural network structures as a specific form of mapping.

**Definition 5 Prime-Flippable Network**. Given an $L + 1$ layer flippable neural network $\mathcal{F} = f_L \circ f_{L-1} \circ \cdots \circ f_1 \circ f_0$, if it becomes non-flippable after removing the last layer $f_L$, i.e., $f_{L-1} \circ \cdots \circ f_1 \circ f_0$ is not flippable, then we call $\mathcal{F}$ a prime-flippable network.

**Theorem 2** Suppose $\mathcal{F}$ is a flippable neural network, disregarding concatenations (for concatenations, refer to Corollary 4 without loss of generality). There must exist a subnet $\mathcal{F}_0$ that is a prime-flippable network such that $\mathcal{F} = g_1 \circ g_2 \circ \cdots \circ g_k \circ \mathcal{F}_0$, where $g_1, g_2, \dots, g_k$ are odd network layers (odd mappings).

**Proof:** If $\mathcal{F}$ has only one layer, then $\mathcal{F}$ is necessarily prime-flippable. Assume $\mathcal{F}$ contains at least two layers, and $\mathcal{F} = g_1 \circ \mathcal{F}_1$. If $\mathcal{F}$ is a prime-flippable network, i.e., $\mathcal{F}_1$ is not flippable, then $\mathcal{F}_0 = \mathcal{F}$. Otherwise, according to Definition 5, $\mathcal{F}_1$ is flippable, and from Lemma 2, it's known that $g_1$ is an odd network layer (odd mapping). Consequently, as we only consider networks with a finite number of layers, a prime-flippable subnet $\mathcal{F}_0$ must exist, and $g_1, g_2, \dots, g_k$ are all odd network layers. ✅

**Proposition 4** Assume $\mathcal{F}$ is a flippable neural network. If its prime-flippable subnet $\mathcal{F}_0$ contains only one neural network layer (excluding activation function layers), then $\mathcal{F}_0(S_1, S_2)$ must be in the form: $W * (S_1 - S_2)$, meaning the weights for $S_1, S_2$ are completely opposite.

**Proof**: First, we denote the input tensor as $S = [S^1, S^2] = \left(S_{i,j,c_{in}}\right)_{i,j=1,..,p;c_{in}=1,2}$, leading to the output tensor from $\mathcal{F}_0$ as $\mathcal{F}_0(S^1, S^2) = \left(O^0_{i,j,c_{\text{out}}}\right)_{i,j=1,\ldots,p;c_{\text{out}}=1,\ldots,C_{\text{OUT}}}$. If it includes only one neural network layer, it can be expressed as: $O^0_{i,j,c_{out}} = \sum_{(u,v)\in\Delta} W_{c_{\text{out}},1,u,v} S_{i+u,j+v,1} + \sum_{(u,v)\in\Delta} W_{c_{out},2,u,v} S_{i+u,j+v,2} + b_{c_{out}}$. Given that $\mathcal{F}_0$ is flippable, we have $\mathcal{F}_0(S^1, S^2) + \mathcal{F}_0(S^2, S^1) = \sum_{(u,v)\in\Delta} \left(W_{c_{\text{out}},1,u,v} + W_{c_{\text{out}},2,u,v}\right)\left(S_{i+u,j+v,1} + S_{i+u,j+v,2}\right) + 2b_{c_{out}} = 0$. Due to the arbitrariness of $i, j, S$, it's inferred that $b_{c_{\text{out}}} = 0$ and $W_{c_{\text{out}},1,u,v} + W_{c_{\text{out}},2,u,v} = 0$, implying $W_{c_{\text{out}},2,u,v} = -W_{c_{\text{out}},1,u,v}$. Therefore, $O^0_{i,j,c_{out}} = \sum_{(u,v)\in\Delta} W_{c_{\text{out}},1,u,v}\left(S_{i+u,j+v,1} - S_{i+u,j+v,2}\right)$, which means $\mathcal{F}_0(S^1, S^2) = W * (S^1 - S^2)$. ✅

#### 2.4 FSDiffNet Properties

**Proposition 5** When $\boldsymbol{p} \geq \mathbf{3}$, combining a diagonal convolution kernel with the 3x3 dilation sequence $\{\mathbf{3}^{\boldsymbol{l}}\}_{\boldsymbol{l}=\mathbf{0},\ldots,\boldsymbol{L}}$, the receptive field of each element in the $\boldsymbol{p} \times \boldsymbol{p}$ matrix $\boldsymbol{Y}$ can cover all elements of the input matrix $\boldsymbol{S}$ using only $\boldsymbol{L} = \lceil \mathbf{log}_{\mathbf{3}}(\boldsymbol{p}) \rceil$ layers.

**Proof:** Consider the element $\boldsymbol{Y}_{\mathbf{1},\boldsymbol{p}}$. Since $\boldsymbol{Y}, \boldsymbol{S}$ are symmetric matrices, the receptive field of $\boldsymbol{Y}$ only needs to cover the upper triangular part of $\boldsymbol{S}$. Moreover, due to the diagonal nature of the DiagConv, the receptive field of $\boldsymbol{Y}_{\mathbf{1},\boldsymbol{p}}$ only needs to cover the upper right quarter of matrix $\boldsymbol{S}$, namely $\boldsymbol{S}_{\mathbf{1}:\frac{\boldsymbol{p}}{\mathbf{2}},\left(\frac{\boldsymbol{p}}{\mathbf{2}}+\mathbf{1}\right):\boldsymbol{p}}$. By combining with the dilation sequence $\{\mathbf{3}^{\boldsymbol{l}}\}_{\boldsymbol{l}=\mathbf{0},\ldots,\boldsymbol{L}}$, the receptive field shape of each element in the non-diagonal parts at layer $\boldsymbol{l}$ is $\mathbf{3}^{\boldsymbol{l}} \times \mathbf{3}^{\boldsymbol{l}}$. In summary, if the receptive field of $\boldsymbol{Y}_{\mathbf{1},\boldsymbol{p}}$ covers all elements of $\boldsymbol{S}$, then the lower left quarter of its $\mathbf{3}^{\boldsymbol{l}} \times \mathbf{3}^{\boldsymbol{l}}$ receptive field needs to cover the upper right quarter of matrix $\boldsymbol{S}$. Therefore, the number of layers $\boldsymbol{L}$ that satisfies this condition follows the equation $\frac{\mathbf{3}^{\boldsymbol{L}}}{\mathbf{2}} = \frac{\boldsymbol{p}}{\mathbf{2}}$, resulting in $\boldsymbol{L} = \lceil \mathbf{log}_{\mathbf{3}}(\boldsymbol{p}) \rceil$. ✅

**Proposition 6** The time complexity of FSDiffNet is $\boldsymbol{O\left(p^2 \log(p)\right)}$.

**Proof:** The computational complexity of FSDiffNet depends on the average kernel size $\boldsymbol{C_{IN} * C_{OUT} * K^2}$ and layer number $\boldsymbol{L}$, where $\boldsymbol{C_{IN}}$ is the number of input channels, $\boldsymbol{C_{OUT}}$ is the number of ouput channels, $\boldsymbol{K}$ is the kernel width. However, according to Proposition 5, as $\boldsymbol{p}$ increases, we can get that $\boldsymbol{L \approx \log_K p}$ is enough to cover all elements in the matrix due to "high dialtion+circular padding" strategy. And usually, we set $\boldsymbol{C_{IN}, C_{OUT}, K}$ as small constants independent with $\boldsymbol{p}$. These lead to a time complexity of $\boldsymbol{O\left(p^2 \log(p)\right)}$. ✅

**Proposition 7** Softsparse, processed Batch Normalization (BN), Diagonal Convolution (DiagConv), and standard bias-free 1x1 convolutions are odd layers.

**Proof:** This can be readily derived from the definitions of these components. ✅

**Theorem 3 FSDiffNet is flippable.**

**Proof:** Consider an FSDiffNet consisting of $\boldsymbol{L}$ layers, whose final output is given by $\boldsymbol{\mathcal{F}^L(S^1, S^2)} = \mathbf{Tanh}\left(\boldsymbol{C}\left(\boldsymbol{\mathcal{F}^{L-1}(S^1, S^2)}\right)\right)$. Starting from the input, trivially, the zeroth layer is defined as $\boldsymbol{\mathcal{F}^0(S^1, S^2) = S^1 - S^2}$, hence $\boldsymbol{\mathcal{F}^0}$ is flippable. The first layer is $\boldsymbol{\mathcal{F}^1(S^1, S^2) = \sigma\left(BN\left(\left[DC(S^1 - S^2), f^1(S^1) - f^1(S^2)\right]\right)\right)}$, where according to Lemma 1, Corollaries 4 and 5, and Proposition 7, $\boldsymbol{\mathcal{F}^1}$ is flippable. For the intermediate layers,

$$\boldsymbol{\mathcal{F}^l(S^1, S^2) = \sigma\left(BN\left(\left[DC\left(\mathcal{F}^{l-1}(S^1, S^2)\right), f^l(S^1) - f^l(S^2)\right]\right)\right)},$$

$$\boldsymbol{\mathcal{F}^{L-1}(S^1, S^2) = BN\left(\left[C\left(\mathcal{F}^{L-2}(S^1, S^2)\right), f^{L-1}(S^1) - f^{l-1}(S^2)\right]\right)},$$

similarly, it can be shown that $\boldsymbol{\mathcal{F}^l}$, $\boldsymbol{\mathcal{F}^{L-1}}$ are flippable. The final output is $\boldsymbol{\mathcal{F}^L(S^1, S^2)} = \mathbf{Tanh}\left(\boldsymbol{C}\left(\boldsymbol{\mathcal{F}^{L-1}(S^1, S^2)}\right)\right)$. Obviously, $\boldsymbol{Tanh}$ is an odd layer. Therefore, according to Corollary 5, $\boldsymbol{\mathcal{F}^L}$ is flippable, implying that the entire FSDiffNet is flippable. ✅

### 3. Supplementary Note 3: Baseline Methods

#### 3.1 Single-condition Baseline

##### 3.1.1 Single-condition architecture

The whole architecture for single-condition graph inference is shown in Figure S1. The major points consist of diagonal convolution kernels, high dilation sequences, circular padding, and an alternative that weakens the diagonal part. The input of this network is the sample covariance matrix $S = cov(X)$ normalized with diagonal 1.

##### 3.1.2 Double condition baseline architecture

In this section, we propose a naïve extension version of the single-condition baseline model, denote as baseline-double (baseline-d). The whole architecture is shown in Figure S2. The only difference with single-condition baseline is that the input is made up of the stack of $S^1, S^2$.

We also introduce another baseline model for further comparison, which is denoted as baseline-single (baseline-s). It is direct subtraction of the output of architecture in 3.1.1, i.e. $f(S^1) - f(S^2)$.

### 4. Supplementary Note 4: Evaluation Metrics

To better compare the performance, we carefully designed metrics for differential inference problems with both positive and negative targets. First, we extend traditional metrics whose labels are $\{\mathbf{0}, \mathbf{1}\}$ to $\{-\mathbf{1}, \mathbf{0}, \mathbf{1}\}$. We separately calculate the positive part and the negative part and finally average these two metrics, i.e., $\boldsymbol{metric} = \frac{\boldsymbol{metric_{pos}} + \boldsymbol{metric_{neg}}}{\mathbf{2}}$ (Figure S3).

### 5. Supplementary Note 5: Training Settings

We use the ‘Adam’ optimizer, ‘ReduceLROnPlateau’ learning schedule.  Generate real-time training sets for each epoch, early stop training if its validation accuracy and loss are not improved. All trainings were completed in parallel on two NVIDIA Quadro P5000 GPUs.

### 6. Supplementary Note 6: Siamese Differential Layer Ablation Experiment

The differential layer is generated by the subtraction of two channels of the siamese network. Intuitively, we aim not only to extract features from the input layer but also to extract features from each differential layer of the siamese network, maximizing the use of information from the siamese network. Siamese networks are traditionally employed to assess the similarity between two inputs, a common application being the authentication of signature veracity. Contrary to this conventional use, our approach utilizes these networks to deduce differences. This methodology represents a notable pivot from qualitative to quantitative analysis.

Specifically, we validated the role of the differential layer through ablation experiments. We first denote the differential layer obtained by subtracting the channels in the middle part of the siamese network as M, and the differential layer obtained by subtracting the output part of the siamese network at the end as E (Figure S5). We then successively remove the differential layers corresponding to M, E, and ME, obtaining FSDiffNet_M, FSDiffNet_E, FSDiffNet_ME respectively. To maintain the scale of the network, the number of yellow channels (trainable part) corresponding to the deleted blue channels is doubled from its original size. It's noted that the initial differential layer is not removed because the yellow channel (trainable part) is derived from the initial differential layer; removing it would render the entire network untrainable. Through experimentation, we observed that removing either M or E would impact the performance of the network inference. Particularly noteworthy is the significant performance decrease observed when both M and E are removed simultaneously (see Figure S6). This experiment confirms that FSDiffNet effectively utilizes differential channel information within the siamese network for differential graph inference.

### 7. Supplementary Note 7: Scale Variability

In the framework of differential network inference based on optimization algorithms, the algorithm does not rely on the scale of input and its performance is usually related to the quality of data. However, methods based on deep learning frameworks are often sensitive to scale, for example, in convolutional neural networks, the size and stride of the convolution kernel can affect the dimensions of the output, and researchers often need to constrain or adjust the scale of input data.

Therefore, in this section, we investigated the scale variability of FSDiffNet. In FSDiffNet, the scale of output remains the same as the input, thus it is not dependent on the number of input variables, $p$. Specifically, we randomly generated training data of different scales, $p \in \{20,40,60,80,100\}, n = 70$ and let FSDiffNet learn from it. Then, we randomly generated test data of different scales to test its performance. First, we fixed $n = 70$ and varied $p \in \{10,20,30 \dots ,90,100\}$, and then observed the changes in performance. As a control, we used the performance of Pinv to indicate the difficulty of the network inference task under this dataset. It can be seen that as $p$ gradually increases, $n/p$ gradually decreases, and the difficulty of the problem gradually increases, the performance of both methods gradually decreases, and the trend is roughly the same (Figure S7). This indicates that the performance loss of FSDiffNet is mainly due to the difficulty of the dataset and not caused by generalization error. Therefore, the FSDiffNet, based on a deep learning architecture, is robust with respect to the dimensionality of variables and has strong generalization capability. Indeed, FSDiffNet can learn effective features for differential network graph from simulated data.

Subsequently, we fixed $p = 40$ and varied $n \in \{10,20,30 \dots ,90,100\}$, and observed the changes in performance. As $n$ gradually increases, $n/p$ gradually increases, and the difficulty of the problem gradually decreases, the AUPR performance of both methods gradually increases (Figure S8). Similarly, we observed the same phenomenon, where the changes in FSDiffNet's performance were mainly due to changes in the complexity of the dataset. This once again confirms that the FSDiffNet we carefully designed exhibits scale variability.

### 8. Supplementary Note 8: Permutability of Variables

In the framework of convolutional neural networks, the permutation of pixels in an input image and rotation of angles can lead to incorrect predictions. Furthermore, scrambling the order of rows and columns in an input image can even render the neural network completely ineffective. However, an interesting feature of FSDiffNet is the permutability of variables, and even integrating swapped results can slightly improve the accuracy of FSDiffNet. Specifically, we conducted the following experiment. First, we randomly generated a permutation matrix $P$, and then obtained the permuted inputs $(PS^1P^T, PS^2P^T)$. Lastly, we used $P^T\mathcal{F}(PS^1P^T, PS^2P^T)P$ as the graph inference result. The result averaged over 100 permutations, denoted as FSDiffNet*100, shows that as the number of permutations increases, both AUPR and Precision@5% slightly increase, indicating that averaging over multiple permutations can to some extent remove part of the noise (Figure S9).

Figure S10 showcases an example of graph inference by FSDiffNet after permutation. It can be observed that after permutation, the main network structure is not lost, but there are slight differences in some of the finer edges. Intuitively, a perturbation can be seen as a new sample within the same sample space, thus it makes sense that FSDiffNet does not lose its capability. In theory, each element in the output matrix is solely influenced by its receptive field, and the network weights remain constant before and after the permutation of input nodes. In conventional convolution methods, the receptive field primarily encompasses elements near the target element, and node permutation can disrupt the neighborhood structure. Therefore, by rapidly expanding the receptive field through diagonal convolution combined with a high dilation rate and circular padding, we can mitigate, to some extent, the effects induced by node permutation.

### 9. Supplementary Note 9: Time Complexity Analysis

We compared the time complexity of each method (Table 1 in the main text). The results show that, as a neural network-based method, FSDiffNet significantly outperforms other methods.

- The time complexity of GLasso varies between $O(p^3) \sim O(p^4)$ depending on the density of the network;
- JGL is related to the number of iterations $T_1$ required for convergence, where each iteration requires at least $O(p^3)$ time complexity;
- BDgraph requires independent calculation of the birth-death rate for each edge, and also needs $T_2$ steps of sampling for the graph and precision matrix. The independent calculation of birth-death rate for each iteration requires $O(p^3)$ complexity, thus totaling $O(T_2p^5)$ time complexity. However, since the birth-death rate for each edge can be calculated in parallel, the complexity can be reduced to $O(T_2p^3)$ when there are enough CPU cores;
- NetDiff conducts regression for each gene separately, thus requiring a total of $O(T_3p^3 * p) = O(T_3p^4)$ time complexity, where $T_3$ is the number of iterations for variational inference. Similarly, $p$ regressions can be computed in parallel, thereby reducing the complexity to $O(T_3p^3)$;
- The complexity of FSDiffNet depends on the size of the convolution kernel $C_{IN} * C_{OUT} * K^2$ and the number of layers $L$. However, according to Proposition 5, as $p$ increases, it can be obtained that $L \approx \log_K p$ layers are sufficient to cover all elements in the matrix. Usually, we set $C_{IN}, C_{OUT}, K$ as constants independent of $p$, for example, in this experiment, $C_{IN} = 50, C_{OUT} = 50, K = 3$.

Another advantage of FSDiffNet is its ability to be easily accelerated using GPUs, a unique advantage of deep learning framework methods.

### 10. Supplementary Figures

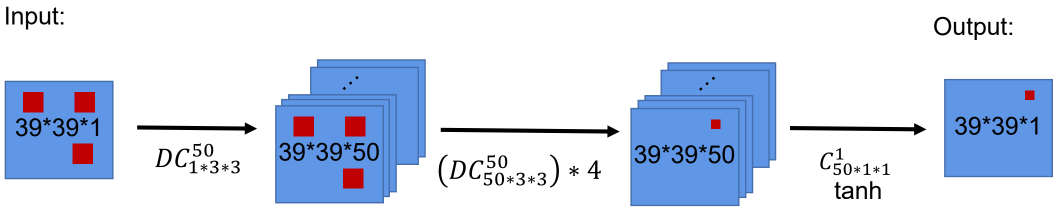

**Figure S1. The single-condition architecture for single-condition graph inference.** DC represents diagonal convolution.

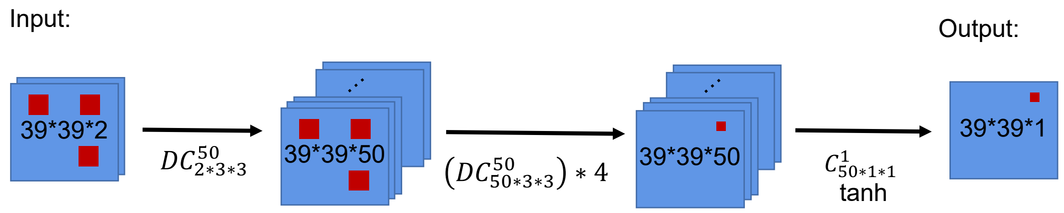

**Figure S2. The architecture of baseline-double (baseline-d).** The only difference with single-condition baseline is that the input is made up of the stack of $\boldsymbol{S}^{1}, \boldsymbol{S}^{2}$.

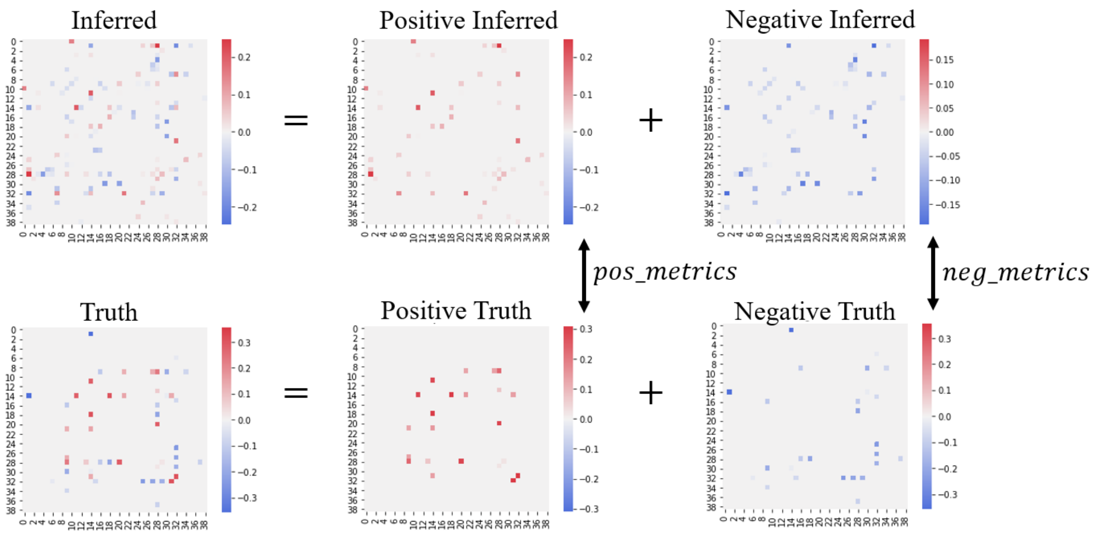

**Figure S3. Diagram of extension metrics.** The metrics are computed separately for positive and negative interactions, and then averaged together.

**Algorithm 1** Simulation data generation algorithm

**Input:**

- $N$: The total number of simulations.
- $\rho$: Sparsity for $\Theta$ matrices.
- $\delta$: Sparsity for $\Delta$ matrices.
- $P_{\text{Dis}}$: The distribution from which $X^{ki}$ samples are drawn.

**Output:**

- $\Theta^{k1}$ and $\Theta^{k2}$: Precision matrices for each simulation $k$.
- $X^{ki}$: Samples generated from the distribution $P_{\text{Dis}}$ based on $\Theta^{ki}$.
- $S^{ki}$: Correlation matrices corresponding to $X^{ki}$.
- $\Delta^k$: The differential matrices between the partial correlation matrices derived from $\Theta^{k1}$ and $\Theta^{k2}$.
- $Y^k$: The sign of $\Delta^k$.

**for** $k = 1$ to $N$ **do**
  *# Generate* $\Theta^{k1}$
  Generate symmetric ER or BA graph weight matrix $\Theta^{k1}$, of whose entry is sampled from $U(-1, 1)$ with sparsity $\rho$;
  *# Generate* $\Theta^{k2}$
  Generate symmetric graph weight matrix $\Delta^{k0}$, of whose entry is sampled from $U(-1, 1)$ with sparsity $\delta$;
  $\Theta^{k2} = \min(\max(\Theta^{k1} + \Delta^{k0}, -1), 1)$
  *# Make* $\Theta^{k1}$, $\Theta^{k2}$ *definite*
  Calculate the min eigenvalues, $e^k = \min\{\text{eig}(\Theta^{k1}), \text{eig}(\Theta^{k2})\}$;
  Add the correction term $|e^k|$ on the diagonal, $\Theta^{ki} = \Theta^{ki} + |e^k|I$, for $i = 1, 2$
  *# Generate simulation data* $X^{ki}$, $S^{ki}$, $\Delta^k$, $Y^k$
  $X^{ki} \sim P_{\text{Dis}}(\Theta^{ki})$, for $i = 1, 2$;
  $S^{ki} = \text{corr}(X^{ki})$;
  $\Delta^k = \text{partial}(\Theta^{k1}) - \text{partial}(\Theta^{k2})$;
  $Y^k = \text{sign}(\Delta^k)$.
**end for**

**Figure S4. Simulation data generation algorithm.**

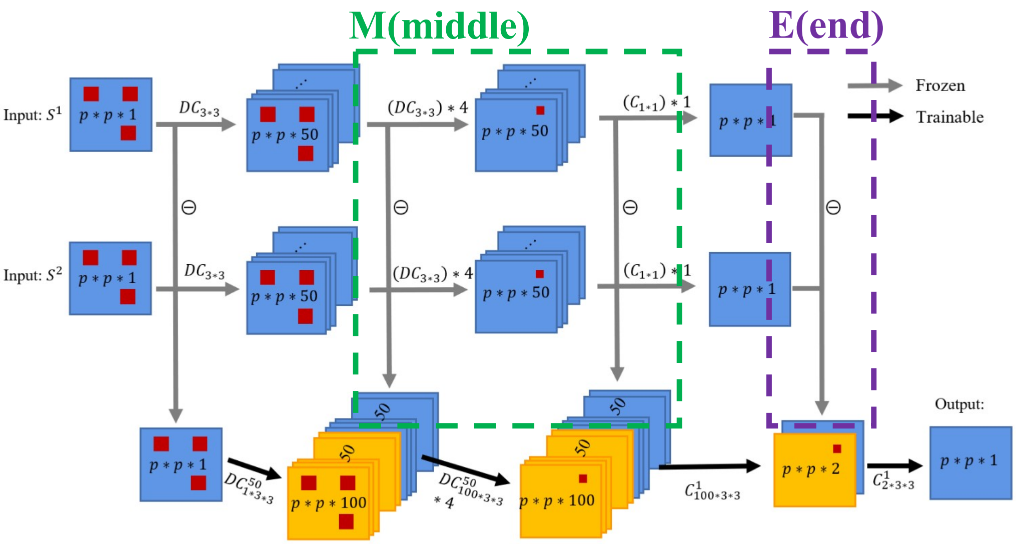

**Figure S5 Schematic diagram of the discrepancy layer ablation experiment.** M represents the middle differential layer and E represents the end differential layer.

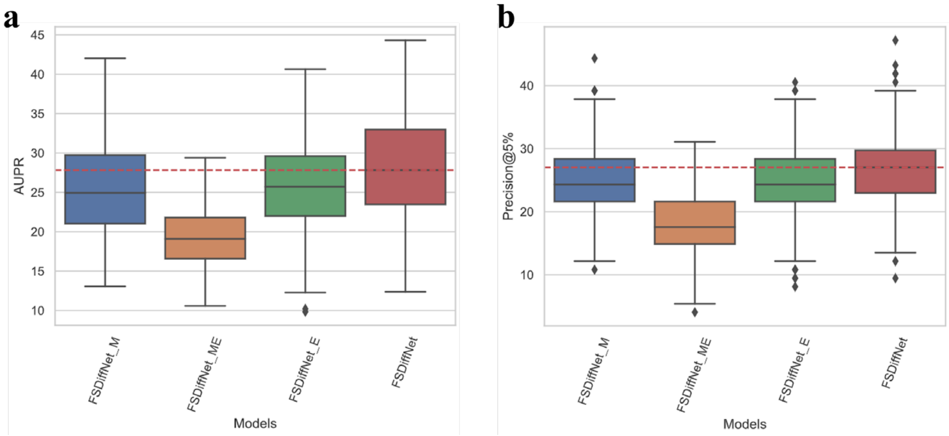

**Figure S6 Comparison of the performance of the ablation experiments.** a) Box line plot for AUPR; b) Box line plot for Precision@5%.

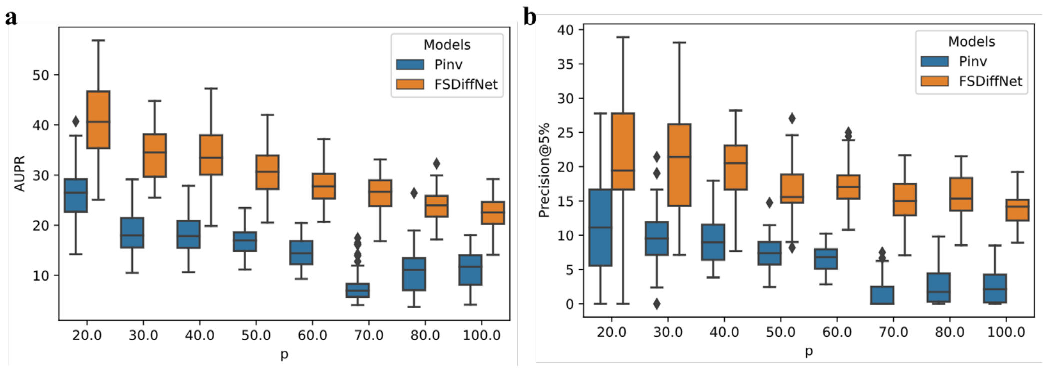

**Figure S7 Variation of FSDiffNet performance with *p***. Pinv is used as a control method to represent the difficulty of the dataset. a) AUPR; b) Precision@5%.

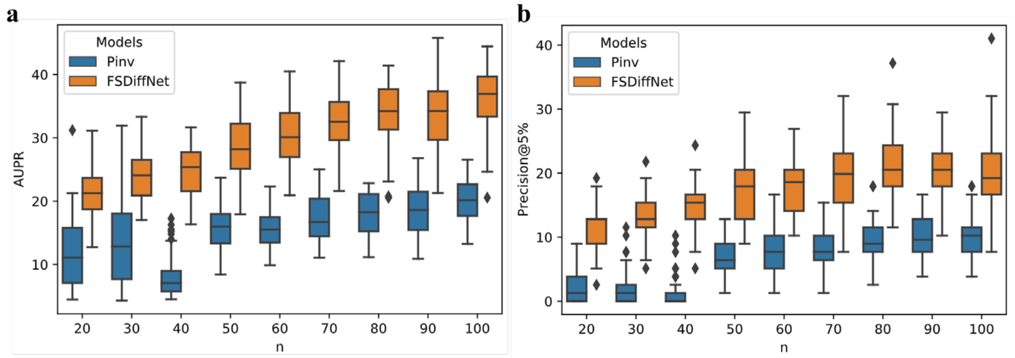

**Figure S8 Variation of FSDiffNet performance with $n$.** Pinv is used as a control method to represent the difficulty of the dataset. a) AUPR; b) Precision@5%

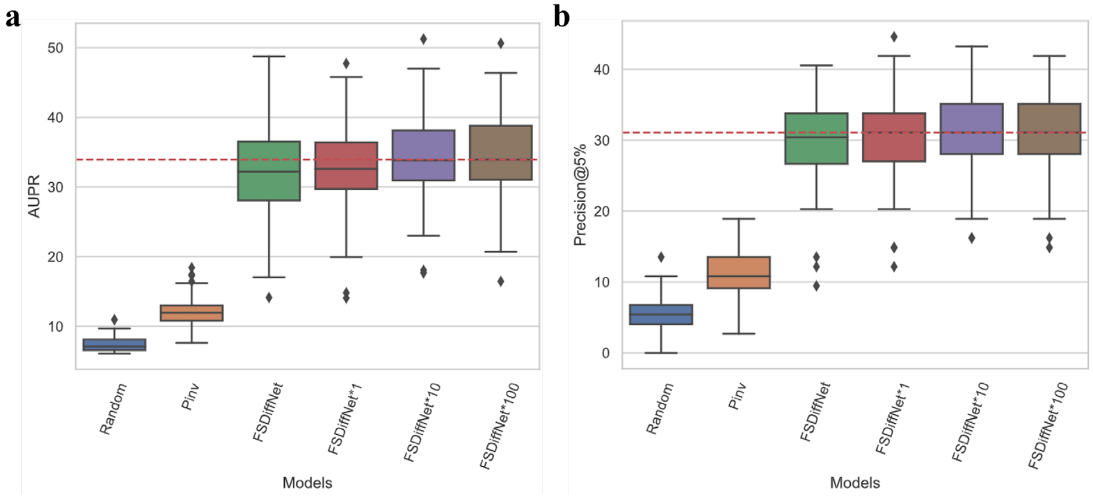

**Figure S9 Comparison of experimental performance box plots for variable permutability.** Pinv is the pseudo-inverse matrix and Random is the random matrix. FSDiffNet****k*** is the average of ***k*** permutation results.

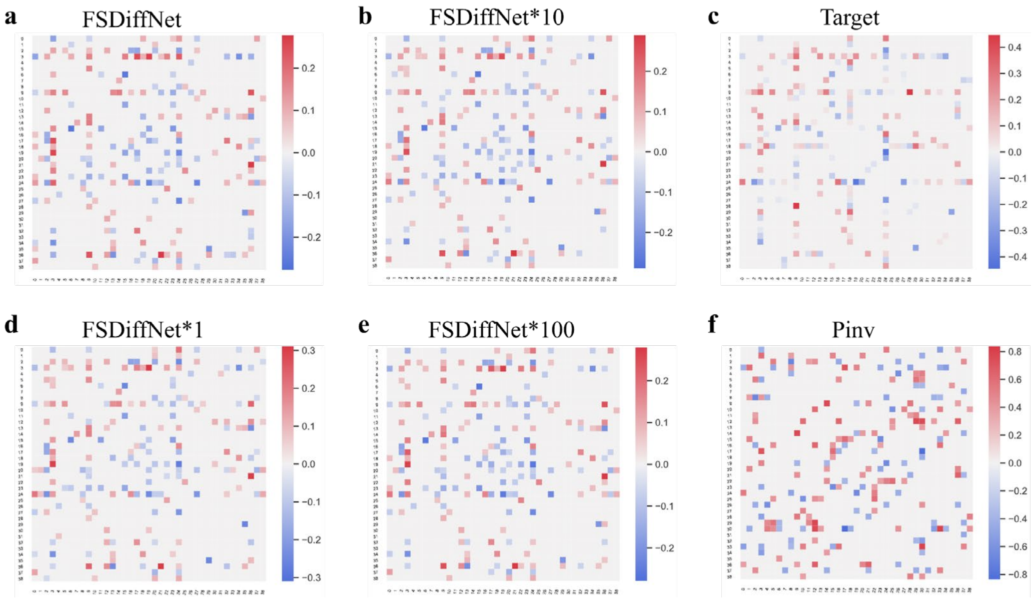

**Figure S10 Example of permutation experiments for 39 variables.** a) FSDiffNet single inference results; b) FSDiffNet permutation 10 times average inference results; c) true labels; d) FSDiffNet permutation 1 time inference results; e) FSDiffNet permutation 100 times average inference results; f) Pinv, inference results of pseudo-inverse method.

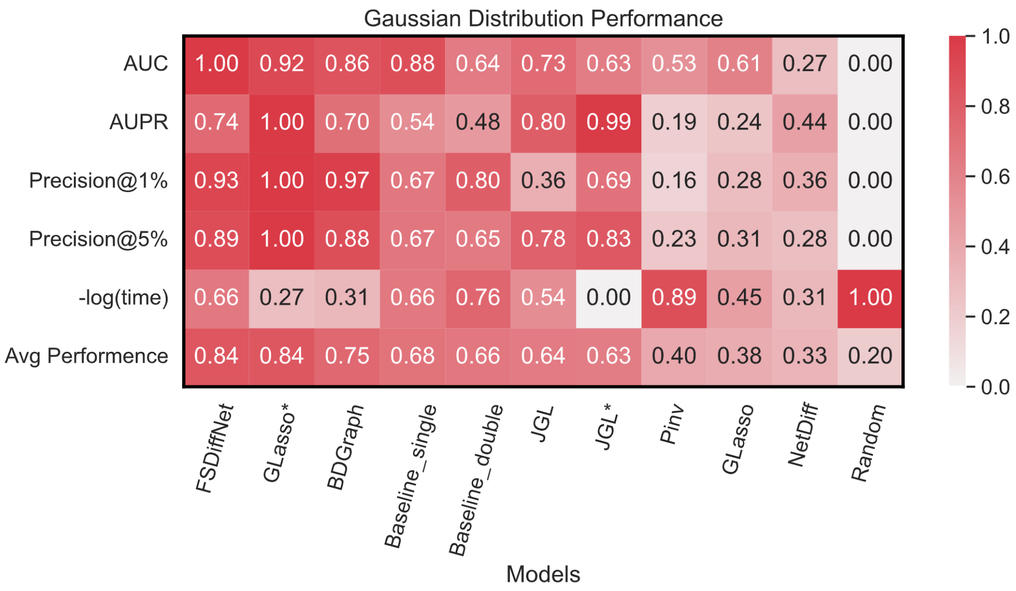

**Figure S11 Heatmap of comprehensive performance comparison under normal scale and gaussian distribution settings.** The vertical axis representing different performance metrics and the last column representing average performance. The horizontal axis lists the different methods, arranged in descending order of average performance, with the highest score being 1 and the lowest being 0.

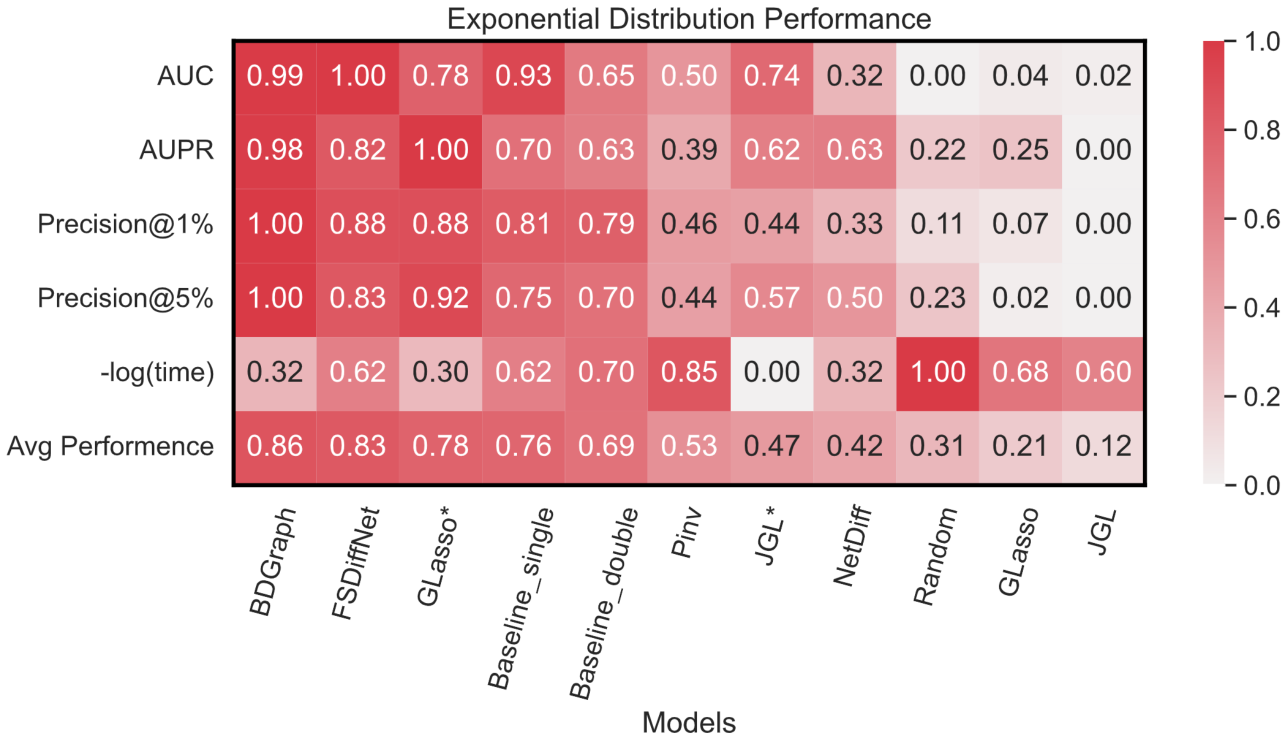

**Figure S12 Heatmap of comprehensive performance comparison under normal scale and exponential distribution settings.** The vertical axis representing different performance metrics and the last column representing average performance. The horizontal axis lists the different methods, arranged in descending order of average performance, with the highest score being 1 and the lowest being 0.
